# A small molecule antagonist for the Tudor domain of SMN disrupts the interaction between SMN and RNAP II

**DOI:** 10.1101/829689

**Authors:** Yanli Liu, Aman Iqbal, Weiguo Li, Zuyao Ni, Yalong Wang, Jurupula Ramprasad, Karan Joshua Abraham, Mengmeng Zhang, Dorothy Yanling Zhao, Su Qin, Peter Loppnau, Xinghua Guo, Mengqi Zhou, Peter J Brown, Xuechu Zhen, Guoqiang Xu, Karim Mekhail, Xingyue Ji, Mark T. Bedford, Jack F. Greenblatt, Jinrong Min

**Author notes:** These authors contributed equally to this work (Yanli Liu, Aman Iqbal, Weiguo Li, Zuyao Ni, Yalong Wang).

## Abstract

Survival of motor neuron (SMN), a Tudor-domain-containing protein, plays an important role in diverse biological pathways via recognition of symmetrically dimethylated arginine (Rme2s) on proteins by its Tudor domain, and deficiency of SMN leads to the motor neuron degenerative disease spinal muscular atrophy (SMA). Here we report a potent and selective antagonist with a 4-iminopyridine scaffold targeting the Tudor domain of SMN. Our structural and mutagenesis studies indicate that the sandwich stacking interactions of SMN and compound **1** play a critical role in selective binding to SMN. Various on-target engagement assays support that compound **1** recognizes SMN specifically in a cellular context. In cell studies display that the SMN antagonist prevent the interaction of SMN with R1810me2s of DNA-directed RNA polymerase II subunit POLR2A and results in transcription termination and R-loop accumulation, mimicking depletion of *SMN*. Thus, in addition to the antisense, RNAi and CRISPR/Cas9 techniques, the potent SMN antagonist could be used as an efficient tool in understanding the biological functions of SMN and molecular etiology in SMA.

---

Survival of motor neuron (SMN) is a core component of the SMN complex, which is essential for biogenesis of small nuclear ribonucleoproteins (snRNPs) by assembling the heptameric Sm ring onto spliceosomal snRNA^1^. The Tudor domain of SMN (Fig. 1a) binds to arginine symmetric-dimethylated (Rme2s) Sm proteins, and this interaction plays a critical role in snRNP assembly^2, 3^. Considering the importance of SMN in the fundamental process of snRNP assembly, it is not surprising that complete loss of SMN is lethal. The human genome contains 2 genes, *SMN1* and *SMN2*, which produce the identical SMN protein. Homozygous deletion or mutation of *SMN1* coupled with a single nucleotide substitution at position 6 of exon 7 (C6T) of *SMN2* is responsible for spinal muscular atrophy (SMA)^4^, the most common genetic cause of infant death with a frequency of 1 in ∼10,000 births^5^.

**Fig. 1.**
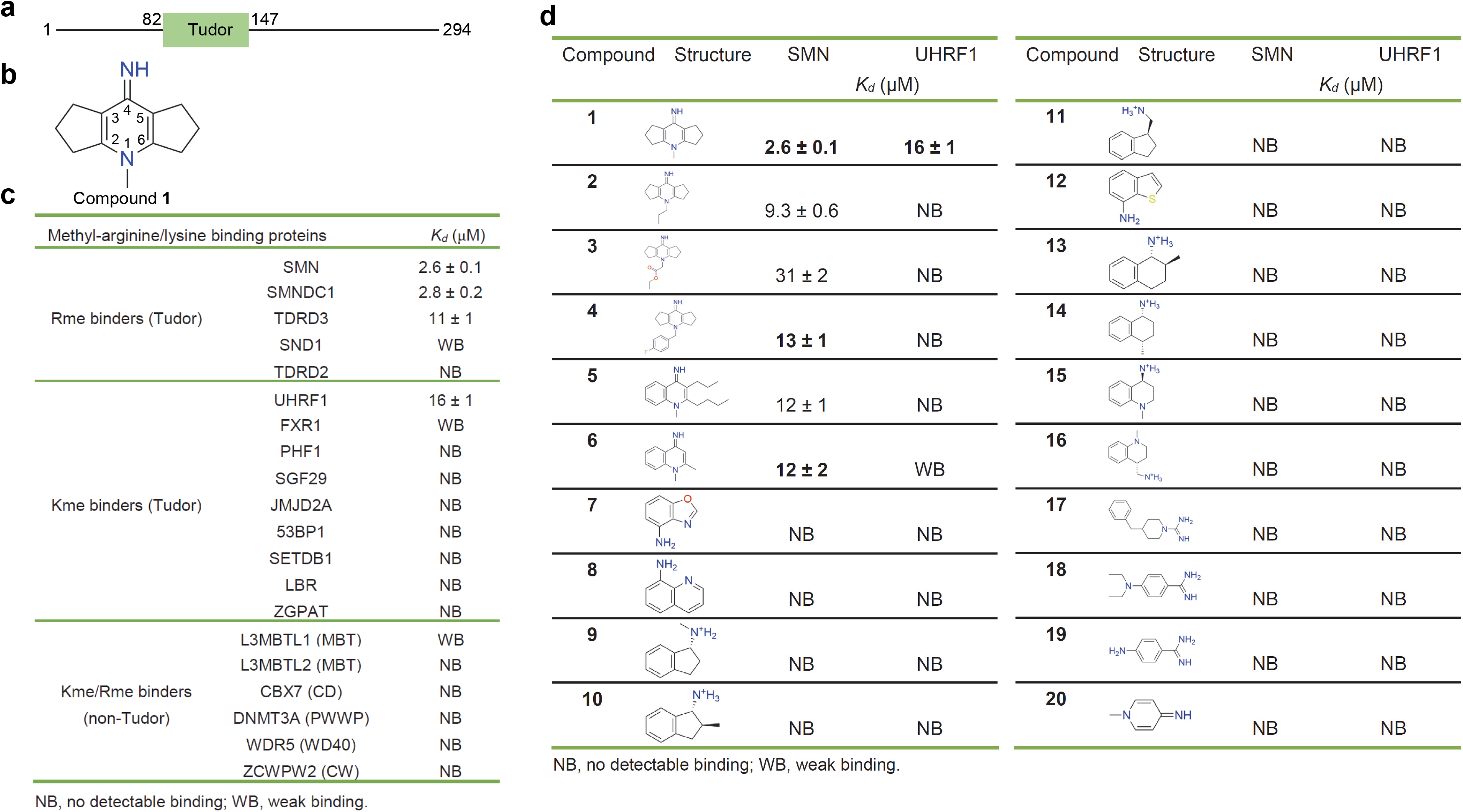
Compound 1 preferentially binds to SMN among assayed methylarginine or methyllysine binders. **a**, Domain structure of SMN. **b**, Molecular structure of compound **1**. **c**, Binding affinities of compound **1** to selected modified histone readers measured by ITC. **d**, Binding affinities of compound **1** analogs reveal the importance of the triple-ring and imino group of compound **1**. ITC data shown are representative of two independent experiments in **c-d**.

In addition to its role in snRNP assembly, SMN is also involved in regulation of nuclear architecture^6, 7^, local axonal translation in neurons^8^ and transcription termination^9^. SMN regulates nuclear architecture by interacting with arginine methylated coilin, a Cajal body (CB) specific protein. Cajal bodies (CBs) and gemini of Cajal bodies (Gems) are the twin sub-cellular organelles in the nucleus of proliferative cells such as embryonic cells, or metabolically active cells such as motor neurons. Coilin harbors symmetrically dimethylated arginine residues^6, 7^. Sufficient arginine methylation of coilin is required for its binding to SMN, which is stored in Gems and accompanies snRNP to CBs during differentiation of the human neuroblastoma cell line SH-SY5Y^10^. SMN is also reported to regulate local axonal translation via the miR-183/mTOR pathway in neurons^8^. Specifically, the miR-183 level is increased and local axonal translation of mTor is reduced in SMN-deficient neurons. In an SMA mouse model, suppression of the miR-183 expression in the spinal motor neurons strengthens motor function and increases survival of *Smn*-mutant mice, which uncovers another potential mechanism responsible for SMA pathology^8^. SMN also interacts with symmetric-dimethylated R1810 at the C-terminal domain (CTD) of RNA polymerase II (RNAP II) subunit POLR2A (R1810me2s-POLR2A) via its Tudor domain to regulate transcription termination^9^. In SMA patients, abnormal transcription termination such as pause of RNAP II and R-loop (DNA-RNA hybrids) accumulation in the termination region may facilitate neurodegeneration^9^. Taken together, SMN functions in different biological pathways, and the Tudor domain of SMN plays a critical role in executing these functions by mediating arginine methylation dependent interactions.

In spite of the extensive study of SMN and its associated SMA disease, it is still unclear how SMN protects motor neurons in the spinal cord against degeneration. To this end, we set out to design SMN-selective chemical probes that would specifically occupy the methylarginine binding pocket and disrupt the Tudor domain-mediated and arginine methylation-dependent interactions. These SMN-specific chemical probes could be used to better understand SMN’s biological functions in different pathways and molecular etiology in SMA.

## Results

### Identification of an SMN-selective small molecule antagonist

In this study, we obtained an SMN-selective antagonist by serendipity when we tried to screen inhibitors against the histone H3K9me3 binding tandem Tudor domain (TTD) of UHRF1 (Fig. 1b and Extended Data Fig. 1a). In this fluorescence-based peptide displacement screen for UHRF1, we found 5 hits, among which compound **1** was confirmed by isothermal titration calorimetry (ITC) (*K_d_* ∼16 µM, Extended Data Fig. 1b and 1c). As we know, many proteins bind to lysine and arginine methylated histones/proteins, including the Tudor Royal superfamily (Tudor, chromodomain, PWWP and MBT) of proteins and some CW and PHD domain containing proteins^11–14^, and all of these proteins utilize an aromatic cage to recognize the methyllysine or methylarginine residue. In order to investigate the binding selectivity of compound **1**, we screened it against selected methylarginine or methyllysine-binding Tudor domains and methyllysine/methylarginine-binding non-Tudor domains (Fig. 1c). UHRF1_TTD was the only assayed methyllysine binder that measurably bound to compound **1**, and compound **1** bound more tightly to the methylarginine binding Tudor domains of SMN, SMNDC1, and TDRD3 than to the methyllysine binding Tudor domain of UHRF1_TTD (Fig. 1c and Extended Data Fig. 2). SMN, SMNDC1 and TDRD3 are the only three known methylarginine binding canonical single Tudor proteins. Moreover, the highly homologous Tudor domains of SMN and SMNDC1 bound to compound **1** with a ∼4-fold selectivity over that of TDRD3 (Fig. 1c).

### Compound 1 specifically engages SMN in a cellular context

To verify the cellular on-target engagement of compound **1**, we subcloned *SMN* into the mammalian expression vector mCherry2-C1 to express the N-terminally mCherry tagged SMN fluorescent protein, and conjugated compound **1** to 9-(2-carboxy-2-cyanovinyl)julolidine (CCVJ), a fluorescent molecular rotor as previously reported^15^ to afford CCVJ-Cmpd **1**, which presents switched-on fluorescence upon binding to SMN, or to biotin to generate a biotin conjugate compound **1** (biotin-Cmpd **1**) for cellular lysate pulldown assays^16^ (Fig. 2a). Our ITC results showed that these two modified compounds still bound to SMN (Extended Data Fig. 3). When the fluorescence-switching CCVJ-Cmpd **1** binds to SMN, restriction of the fluorescent molecule rotations would trigger emission of strong green fluorescence signals^15^, which is confirmed in solution (Extended Data Fig. 4). Upon treatment of U2OS cells with CCVJ-Cmpd **1**, the green fluorescent compound **1** colocalized with the red fluorescent mCherry-SMN, which was not observed with the SMN cage mutant (Fig. 2b), indicating that compound **1** binds to the aromatic cage of SMN specifically.

**Fig. 2.**
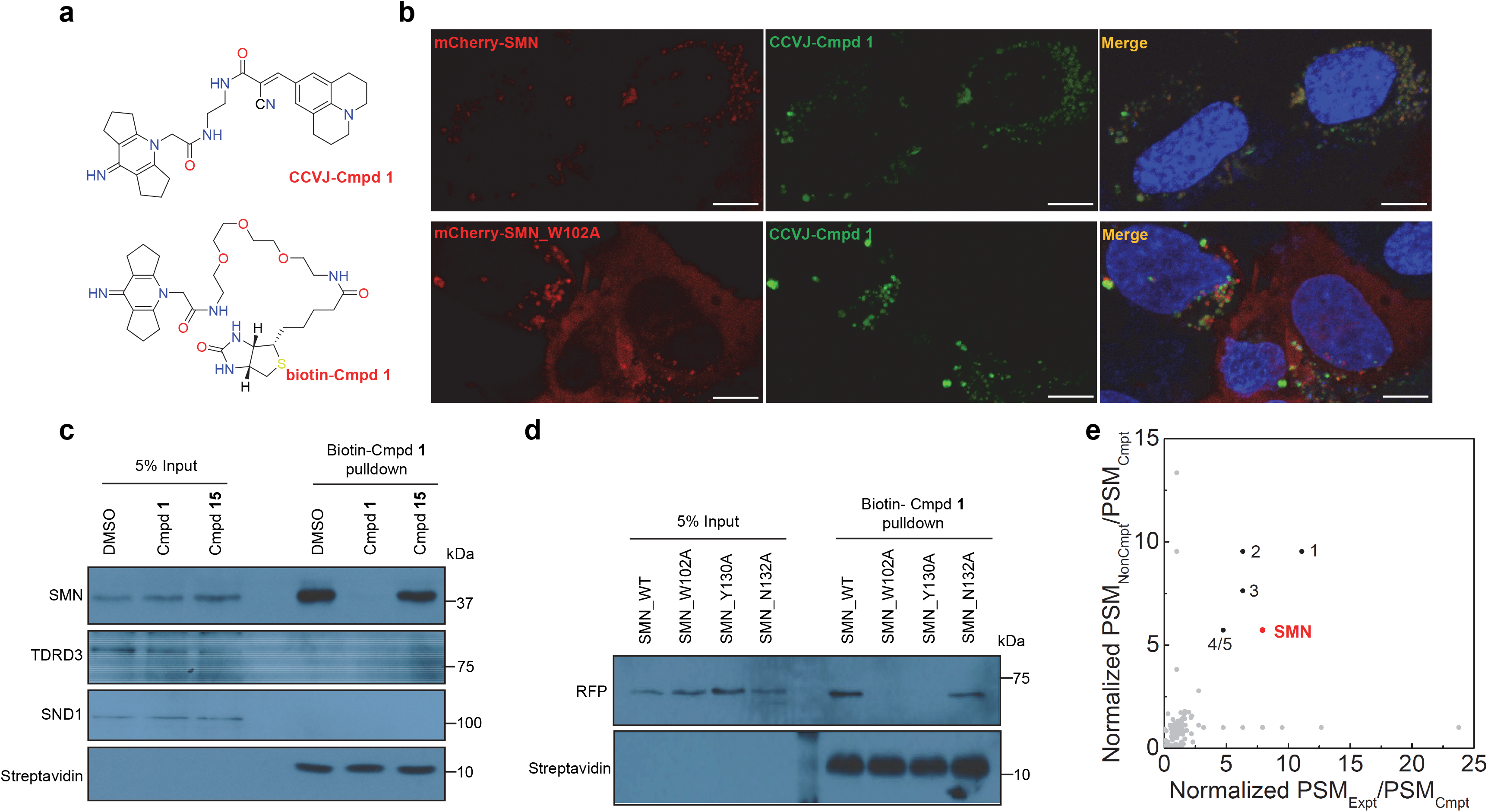
Cellular on-target engagement of compound 1. **a**, Chemical structure of CCVJ conjugated compound **1** (CCVJ-Cmpd **1**) and biotin conjugated compound **1** (biotin-Cmpd **1**). **b**, mCherry-SMN (red) colocalizes with CCVJ-Cmpd **1** (green), which is lost when the cage residue W102 is mutated in SMN. U2OS cells were treated with CCVJ-Cmpd **1** (a final concentration of 10 μM) for 24 h after transfection with 1.0 μg of mCherry-SMN WT (wild-type) or mutant plasmids. Scale bar is 10 μm. Shown are representative confocal images of three independent experiments. **c**, Compound **1**, but not negative compound **15**, inhibits pulldown of SMN by biotin-labeled compound **1** from cell lysates. **d**, SMN cage mutants disrupt or weaken the interaction between SMN and biotin-Cmpd **1** in cell lysates. The U2OS cell lysate was incubated with 20 μM of biotin-Cmpd **1** overnight at 4 °C in figures c and d. For the competition analysis, 100 μM of biotin-free compound **1** or compound **15** was added at the same time when the biotin-Cmpd **1** was incubated with the cell lysates in figure **c**. Data shown are representative of three independent experiments in **c-d**. **e**, On-target engagement of compound **1** is analyzed by chemical proteomics. The PSMs of MS data of 20 μM biotin-Cmpd **1** (Expt) and 20 μM biotin-Cmpd **1** together with 100 μM negative (non-competitive) compound **15** (biotin-free, NonCmpt) were normalized by MS data of 20 μM biotin-Cmpd **1** together with 100 μM competitive compound **1** (biotin-free, Cmpt) to obtain normalized PSM_Expt_/PSM_Cmpt_ and normalized PSM_NonCmpt_/PSM_Cmpt_, respectively, which were used to prepare figure by Origin software 7.0. 1, MOV10 (RNA helicase MOV-10), 2, DKC1 (H/ACA ribonucleoprotein complex subunit DKC1), 3, RPL5 (60S ribosomal protein L5), 4/5, RPL18 (60S ribosomal protein L18) and SSB (Lupus La protein).

To further confirm the cellular binding of compound **1** to SMN, we performed pulldown assays of cell lysates by using the biotin-labeled compound **1** (biotin-Cmpd **1**). The results showed that SMN could be efficiently captured, while neither TDRD3 nor SND1 could be detected (Fig. 2c and Extended Data Fig. 5). The biotin-Cmpd **1** could be competed out by the presence of unlabeled compound **1,** but not the negative control compound **15** in the lysates (Fig. 2c). Furthermore, the biotin-Cmpd **1** could not efficiently pulldown the SMN cage mutants (Fig. 2d). Affinity-purification and mass spectrometry (AP-MS) based proteomic analysis of the biotin-Cmpd **1** pulldown samples identified SMN as one of the 6 significant protein targets. Their interaction was efficiently blocked by the competition of compound **1**, but not by the negative control compound **15** (Fig. 2e). The other AP-MS prey proteins, such as MOV10 (RNA helicase MOV-10), DKC1 (H/ACA ribonucleoprotein complex subunit DKC1), RPL5 (60S ribosomal protein L5), RPL18 (60S ribosomal protein L18) and SSB (Lupus La protein) are common contaminant proteins found in streptavidin affinity-purified samples from the U2OS cell lysates (CRAPome, Contaminant Repository for Affinity Purification, http://crapome.org/). In addition, the AP-MS data from the GFP-SMN overexpressed U2OS cell lysate also showed that SMN was enriched specifically by the biotin-Cmpd **1**, accompanied by two other common contaminants of the U2OS cell lysate, KRT8 (Keratin, type II cytoskeletal 8) and HNRNPL (Heterogeneous nuclear ribonucleoprotein L) (Extended Data Fig. 6). Taken together, these data provide convincing evidence that compound **1** binds to the full-length SMN protein specifically in a cellular context.

### Structural basis of selective compound 1 binding to SMN

In order to understand the structural basis of compound **1** recognition by these reader proteins, we determined the crystal structures of compound **1** in complex with SMN, TDRD3 and UHRF1, respectively (Fig. 3, Extended Data Figs. 7/8 and Table 1). In the complex structure of SMN-compound **1**, compound **1** bound to the aromatic cage formed by W102, Y109, Y127 and Y130, which otherwise accommodates dimethylarginine of its physiological ligands (Fig. 3a-3c). W102 and Y130 sandwich compound **1** rings. In addition, compound **1** forms a hydrogen bond between its imino group and the side chain of N132. This hydrogen bond boosts the ligand binding ability of SMN because mutating N132 to alanine significantly reduced its binding affinity (Fig. 3d).

**Fig. 3.**
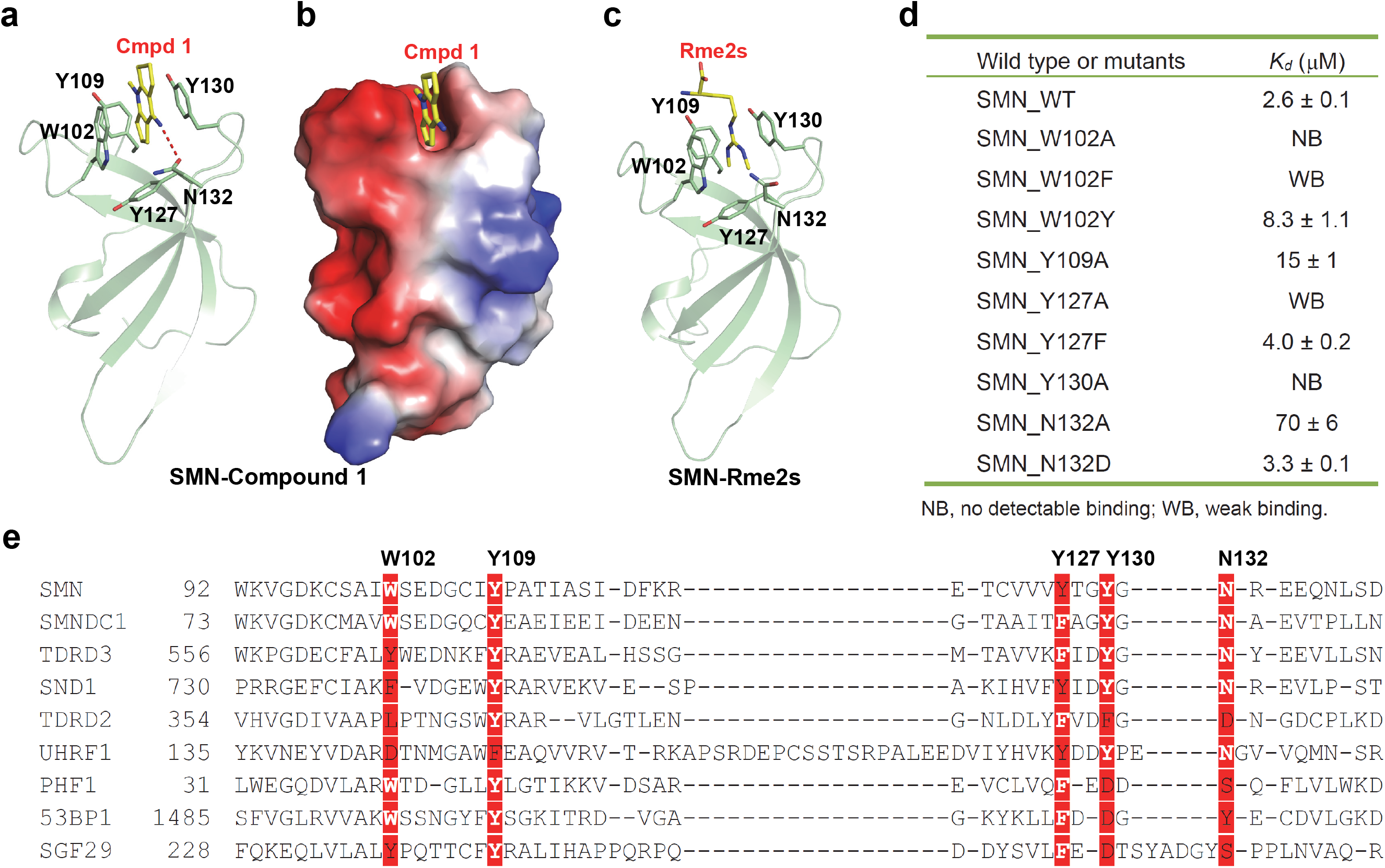
Structural basis of preferential binding of compound 1 to SMN. **a**, Complex structure of Tudor domain of SMN and compound **1** is shown in a cartoon mode. The Tudor domain of SMN is colored in green, with the interacting residues shown in sticks and the intermolecular hydrogen bonds indicated by red dashes. **b**, Electrostatic potential surface representation of the complex of Tudor domain of SMN and compound **1**. **c**, Complex structure of Tudor domain of SMN and Rme2s shown in a cartoon mode. **d**, Binding affinities of compound **1** to different SMN Tudor mutants determined by ITC. Shown are representative of two independent experiments. **e**, Sequence alignment of selected Tudor domains. The compound **1** interacting residues are highlighted in red background. Structure figures were generated by using PyMOL. Surface representations were calculated with the built-in protein contact potential function of PyMOL.

**Table 1.**
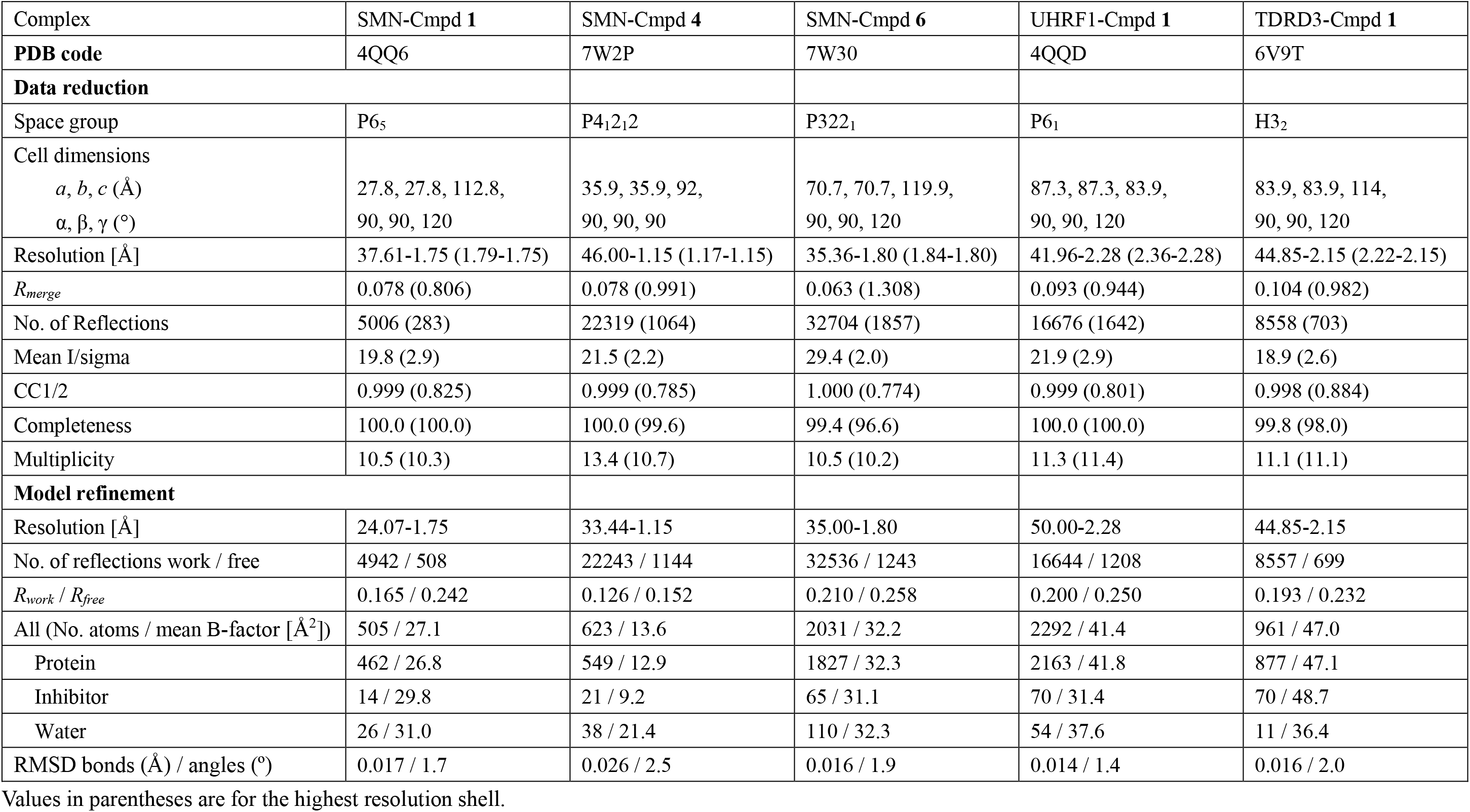
Data collection and refinement statistics.

In the TDRD3-compound **1** structure, the binding mode is largely conserved (Extended Data Fig. 7), but Y566 of TDRD3 might not stack with the compound as effectively as W102 in SMN (Fig. 3e and Extended Data Fig. 9), which may explain weaker affinity of TDRD3 to compound **1** (Fig. 1c). Consistent with this hypothesis, the W102Y mutant of SMN showed a similar binding affinity to compound **1** as TRDR3 (Fig. 3d). Although we did not determine the corresponding complex structure of SND1, the structure-based sequence alignment revealed that W102 of SMN corresponds to F740 of SND1 (Fig. 3e and Extended Data Fig. 9). Consistent with this, the W102F SMN mutant only weakly bound to compound **1** (Fig. 3d). In addition, the W102A and Y130A SMN mutants did not bind to compound **1** (Fig. 3d). This lack of binding is consistent with our failure to observe binding for the other tested proteins. On the other hand, the cage mutants Y109A and Y127A of SMN weakened, but did not abrogate their binding to compound **1** (Fig. 3d). Hence, the sandwich stacking interactions of compound **1** by W102 and Y130 play a more critical role in the compound **1** recognition.

### UHRF1 recognizes compound 1 via an arginine-binding pocket, but not the methyllysine-binding aromatic cage

To uncover the specific interactions between UHRF1_TTD and compound **1**, we also solved the complex structure of UHRF1_TTD-compound **1** (Extended Data Fig. 8). Two UHRF1_TTD molecules are present in each asymmetric unit of the UHRF1-compound **1** complex structure (Extended Data Fig. 8a), but we only observed the expected disc-shaped electron density of compound **1** in the histone H3K9me3-binding cage of one UHRF1_TTD molecule, while we found a differently shaped blob in the histone H3K9me3-binding cage of the other UHRF1_TTD molecule (Extended Data Fig. 8b). In the complex structure of UHRF1_TTD-PHD and the H3K9me3 peptide (PDB code: 3ASK), an arginine residue R296 in the linker between the TTD and PHD domains of UHRF1 is found in a pocket formed by D142, E153, A208, M224, W238 and F278 from the TTD domain^17^ (Extended Data Figs. 1a and 8c). R296 is fixed in the pocket by forming a salt bridge with D142. Intriguingly, we found the disc-shaped density that resembles the compound **1** in the arginine-binding pockets of both UHRF1_TTD molecules, and compound **1** is stacked between the indole ring system of W238 and the guanidinium group of R209 in both UHRF1_TTD molecules (Extended Data Fig. 8d).

Several lines of evidence suggested that the arginine-binding pocket is the major binding site and the methyllysine-binding aromatic cage is just a minor or non-specific binding site: First, when we mutated the aromatic cage residues of UHRF1_TTD that have been shown to be critical for histone H3K9me3 binding to alanine, the binding affinity of compound **1** is not affected significantly. In contrast, when we mutated the arginine-binding pocket residues, the binding is totally disrupted (Extended Data Fig. 8e). Second, the electron density inside the H3K9me3 aromatic cage is either smear and can be modelled in multiple orientations of compound **1**, which implies that compound **1** does not bind to the cage specifically, or is of no defined density shape (Extended Data Fig. 8b). Third, the aromatic cage has a propensity to accommodate small molecules non-specifically. For instance, some molecules in buffer have been found in the aromatic cage of TDRD3^18^. For the case of UHRF1_TTD, some ethylene glycol molecules from the crystallization buffer are found in the H3K9me3 aromatic cage and the arginine-binding pocket of the apo-UHRF1_TTD structure^19^ (Extended Data Fig. 8f). Fourth, in the SMN-compound **1** complex structure, compound **1** is stacked between the aromatic rings of W102 and Y130. However, the three aromatic residues in the aromatic cage of UHRF1 are perpendicular to each other, which could not stack the compound like SMN does (Extended Data Fig. 8g). Taken together, UHRF1 used the arginine-binding pocket to specifically bind to compound **1**, and this arginine-binding pocket could serve as a novel therapeutic venue for designing potent small molecule allosteric regulators of the UHRF1 functions. Based on the structural information we obtained for our UHRF1_TTD-compound **1** complex and the UHRF1_TTD-PHD-H3K9me3 complex (PDB code: 3ASK), it is conceivable that the compound **1** binding pocket of UHRF1 is occupied by R296 of the full-length UHRF1 protein, which would prevent compound **1** from chemiprecipitating UHRF1 from the U2OS cell lysate.

### The imino group of compound 1 plays a critical role in binding to SMN

To explore structure-activity relationship between SMN and compound **1**-like compounds, we procured commercially available analogs that include single, double and triple ring molecules and measured their affinities toward SMN and UHRF1, respectively (Fig. 1d). SMN bound to all the four triple-ring compounds with affinities between 2.6 μM to 31 μM (Fig. 1d and Extended Data Fig. 10). The ligands have a 4-iminopyridine scaffold in common, and none of them is more potent than the original hit. Although the binding affinities of SMN and these compounds are not high, SMN binds to these compounds much stronger than its physiological ligands such as symmetric dimethylarginine or R1810me2s-POLR2A, which showed a binding affinity of 476 μM^20^ or 175 μM^9^, respectively.

Due to the electronic similarity between the N-methyl and 4-imino sites on the 4-iminopyridine core, we did not expect to resolve the orientation of compound **1** based on electron density alone, and could not exclude the possibility that the imino group would instead protrude into the solvent. However, the positive binding results of 1-substituted pyridine cores to SMN presented here could confirm that N132 does interact with the imino group and substituents on the pyridine nitrogen would point away from the Tudor domain, as larger 1-substituents would otherwise clash inside the aromatic cage. In addition, the modifications at the N-methyl site of compound **1**, such as CCVJ-Cmpd **1** and biotin-Cmpd **1**, did not disrupt the interaction of SMN and the modified compounds, which further validates our statement that SMN binds to the 4-imino group of compound **1** to entail a hydrogen bond between the 4-imino group and N132. Indeed, our crystal structure of SMN in complex with compound **4** also confirms that the imino group of compound **4** forms a hydrogen bond with N132 (Extended Data Fig. 11a-b).

In addition to triple-ring compounds **1** to **4**, SMN also bound to two of twelve double-ring compounds, compounds **5** and **6** (Fig. 1d and Extended Data Fig. 10). Both compounds retain the imino group, which pinpoints the importance of the imino group-mediated hydrogen bond in the compound binding and is consistent with our crystal structures of SMN in complex with compounds **1**, **4**, and **6** (Fig. 3 and Extended Data Fig. 11). None of the single-ring compounds bound to SMN, which may not be able to provide strong enough π-π stacking interaction to hold the compounds. UHRF1, however, did not bind to any other three-ring compounds, because the substituents on the pyridine nitrogen of these compounds are too large for the more enclosed arginine-binding pocket of UHRF1.

### SMN antagonists disrupt the SMN-RNAP II interaction

We previously showed that R1810 in the CTD of the mammalian RNAP II subunit POLR2A is symmetrically dimethylated by PRMT5 and the R1810 methylated CTD directly recruits the Tudor-domain protein SMN, which contributes to the assembly of an R-loop resolving complex on the RNAP II CTD^9^. Hence, we asked whether these small molecule antagonists might be able to disrupt the interaction of SMN with RNAP II in cells. To test this possibility, we treated the HEK293 cells with a series of concentrations of either compound **1**, compound **2** or negative control compound **15** for 72 h, and then performed immunoprecipitation using the cell extracts. We demonstrated that coimmunoprecipitation of SMN with GFP fusion protein of POLR2D, a component of RNAP II, was inhibited by compound **1** and compound **2** on a dose-dependent manner, but not by negative compound **15** or DMSO (Fig. 4a). Furthermore, the coimmunoprecipitation of POLR2A with SMN was also disrupted on the treatment of compound **1** and compound **2**, whereas no significant effect was observed for the cells treated with DMSO as a control (Extended Data Fig. 12). These results provide convincing evidence that the small molecule antagonists compound **1** or compound **2** of SMN disrupts the interaction between SMN and the RNAP II complex. Since 20 µM of either compound **1** or compound **2** is enough to exhibit significant inhibition of the interaction between SMN and POLR2A (Extended Data Fig. 12), we used this concentration in the following assays.

**Fig. 4.**
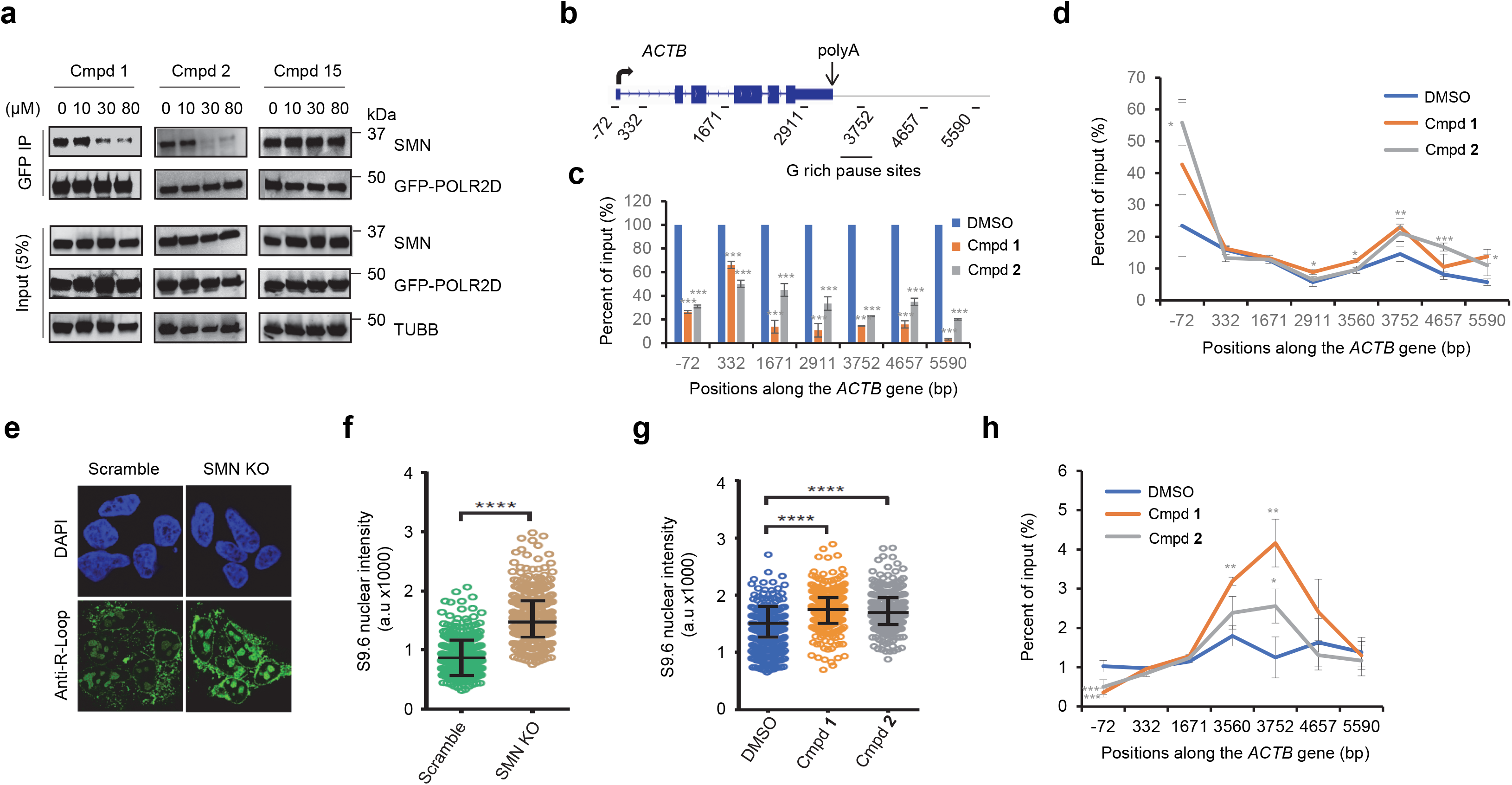
The effects of SMN antagonists on the interaction between SMN and RNAP II, RNAP II pause and R-loop accumulation. **a**, SMN antagonists disrupt binding of SMN to RNAP II. IP-western blotting experiments were performed by using the indicated antibodies in extracts prepared from HEK293 cells expressing GFP-tagged POLR2D in treatment with indicated concentrations of compound **1**, compound **2**, compound **15** or DMSO for 72 h. Data shown are representative of three independent experiments. **b**, Schematic of *ACTB* gene. **c,** SMN antagonists reduce SMN association at *ACTB* gene locus. Quantification of SMN qPCR data from ChIP experiments using SMN antibodies at the indicated *ACTB* primer positions in HEK293 cells. The SMN levels in DMSO controls were set as 100%. **d**, SMN antagonists lead to RNAP II pause. Quantification of RNAP II qPCR data from ChIP experiments using POLR2A antibodies at the indicated *ACTB* primer positions in HEK293 cells. **e**, SMN knockout (KO) leads to R-loop accumulation. Representative single-plane images of Z-stacks of the R-loop levels in scramble *vs* SMN KO cells of three independent experiments. **f**, Global nuclear R-loop accumulation caused by SMN knockout. Scatter plots representing data from single-cell and R-loop immunofluorescence analysis (N = 150 cells per condition; Mean ± Quartiles; ****: *P* < 0.0001 for the Mann-Whitney test). **g**, SMN antagonists cause global nuclear R-loop accumulation. Single-cell and R-loop immunofluorescence analysis of HEK293 cells treated with vehicle (DMSO), 20 μM compound **1** or compound **2** (N = 150 cells per condition; Mean ± Quartiles; ****: *P* < 0.0001 for the Mann-Whitney test). **h**, SMN antagonists lead to R-loop accumulation at *ACTB* gene locus. Quantified DNA immunoprecipitation using primers along *ACTB* locus by using GFP antibodies, in cell extracts that were transfected with GFP-RNase H1 R-loop-binding domain (GFP-HB) fusion construct for R-loop detection. For the figures **c**, **d**, **h**, data are presented as the mean ± S.E.M. of three independent experiments. *: *P* < 0.05, **: *P* < 0.01, ***: *P* < 0.001 for the two-tailed Student’s t-test. For the analysis of the SMN chemical antagonists, HEK293 cells were treated with DMSO, 20 μM of compound **1** or compound **2** for 72 h (figures **c**, **d**, **g**, **h)**. Global nuclear R-loop detection was ascertained via R-loop immunofluorescence using S9.6 antibody for R-loop (Kerafast, ENH001) of the SMN wild-type (scramble), SMN KO and chemical antagonists treated HEK293 cells (figures **e**, **f**, **g**).

### SMN antagonists disrupt SMN gene occupancy and lead to RNAP II pause

Our previous ChIP study has shown that SMN occupies the *ACTN (β-actin)* gene from its promoter to the termination regions with the highest level of occupancy at the 3′-end of the gene^9^. PRMT5 depletion or POLR2A R1810 mutation leads to a decreased SMN occupancy^9^. To examine whether the small molecule antagonists of SMN have any effects on the SMN occupancy at its target genes during transcription, the SMN ChIP assay was performed using the primers along the *ACTN* gene (Fig. 4b). Similar to the effects of PRMT5 depletion or POLR2A R1810 mutation, treatment of either compound **1** or compound **2** significantly reduced the levels of SMN along the *ACTN* gene (Fig. 4c). Given that POLR2A CTD R1810A mutation or depletion of *SMN* leads to the accumulation of RNPA II at genes^9^, the SMN antagonists might have similar effects. To address this issue, we performed RNAP II ChIP experiments and found that addition of either compound **1** or compound **2** (20 µM, 72 h) significantly increased the occupancy levels of RNAP II at the promoter regions and 3′-end of the *ACTB* gene as detected by quantitative PCR (qPCR) (Fig. 4d). These results indicate that the SMN antagonists could cause the accumulation of RNAP II at both the promoter and 3′ pause site of its target genes.

### SMN depletion or its inhibition causes accumulation of the R-loop

Our previous studies demonstrated that PRMT5 or SMN depletion, or POLR2A R1810 mutation leads to the R-loop accumulation at the *ACTB* gene^9^. Here, we further confirmed that the CRISPR/Cas9 mediated SMN knockout increased the global R-loop accumulation in HEK293 cells as detected by immunofluorescence staining (Fig. 4e, 4f and Extended Data Fig. 13). Overexpression of RNase H1 significantly decreased the levels of R-loops in the SMN knockout cells, validating the authenticity of the R-loop signals (Extended Data Fig. 14). Similar to SMN knockout, treatment of either compound **1** or compound **2** significantly increased R-loop levels in comparison to the DMSO controls (Fig. 4g), indicating the global effects of the SMN antagonists in R-loop accumulation. Consistently, treatment of either compound **1** or compound **2** significantly increased the R-loop signals at the 3′-end of the *ACTB* gene (Fig. 4h).

## Discussion

In this study, we identified some low micromolar antagonists with a 4-iminopyridine scaffold targeting the Tudor domain of SMN, and compound **1** shows >4-fold selectivity over other tested methyllysine or methylarginine binding domains. Although the binding affinity of SMN and compound **1** is not high, SMN binds to compound **1** 60–180-fold more tightly than its physiological ligands such as symmetric dimethylarginine or R1810me2s-POLR2A (*K_d_* of 2.6 μM for compound **1** *vs* 476 μM for symmetric dimethylarginine^20^ or 175 μM for R1810me2s-POLR2A^9^). We then utilized different cellular on-target engagement assays to validate that compound **1** specifically recognizes SMN in a cellular context, and showed that compound **1** would prevent the interaction of SMN with R1810me2s of DNA-directed RNA polymerase II subunit POLR2A and result in transcription termination and R-loop accumulation. Hence, compound **1** is a potent and selective antagonist of SMN.

Our structural and mutagenesis studies provide mechanistic insights into the selectivity of compound **1** for SMN. Our protein-compound complex structures uncover that compound **1** is an antagonist targeting methylated arginine binding protein and the sandwich stacking interactions of compound **1** by W102 and Y130 of SMN play a critical role in the compound **1** recognition. The larger binuclear ring structure of tryptophan provides a stronger π-π interaction with compound **1** than tyrosine or phenylalanine. In order to explore if mutating Y130 to tryptophan would increases its binding to compound **1** further, we made a Y130W mutant, which renders the protein to become insoluble, presumably due to steric clash around the aromatic cage. In addition, our structural study uncovers that UHRF1 used an arginine-binding pocket to specifically bind to compound **1**, which indicates that the arginine-binding pocket could serve as a novel therapeutic venue for designing potent small molecule allosteric regulators of the UHRF1 functions.

Although the causative link between SMN deficiency and SMA was established 20 years ago^4^, it remains elusive how deficiency of a protein, which is ubiquitously expressed and causes widespread defects in pre-mRNA splicing in cell culture and mouse models of SMA, would result in a cell-type-specific phenotype: motor neuron dysfunction^21^. A *Drosophila* model suggests that involvement of SMN in snRNP biogenesis does not explain locomotion and viability defects of *Smn* null mutants, implying that SMN may have other functions contributing to the etiology of SMA^22^. Indeed, in addition to its role in snRNP assembly, SMN is also involved in regulation of nuclear architecture by interacting with arginine methylated coilin^6, 7^, local axonal translation in neurons by participating in miR-183/mTOR pathway^8^ and transcription termination by interacting with arginine methylated POLR2A^9^. All of these findings may have important implications for understanding the cell-specific functions of SMN, and shed light on the molecular mechanism of SMA pathology^6–8, 10^. SMN was also proposed to have other functions. It interacts with the mSin3A/HDAC transcription corepressor complex, and represses transcription in an HDAC-dependent manner^23^. In contrast, by interacting with the nuclear transcription activator E2 of papillomavirus, SMN positively regulates E2-dependent transcription^24^. The Tudor domain of SMN recognizes arginine methylated Epstein-Barr virus nuclear antigen 2 (EBNA2), the main viral transactivator of Epstein-Barr virus ^4^, and regulates EBV-mediated B-cell transformation^25^. Infection with the EBV virus can lead to a number of human diseases including Hodgkin’s and Burkitt’s lymphomas. SMN interacts with the fused in sarcoma (FUS) protein, a genetic factor in amyotrophic lateral sclerosis, which links the two motor neuron diseases^26^.

In order to facilitate the elucidation of the biological functions of SMN in different pathways and molecular etiology in SMA, we set out to develop SMN-specific chemical probes, and identified compound **1**, a 2.6 μM antagonist. Although compound **1** could not be claimed as a chemical probe, compound **1** and even weaker binding compound **2** bind to SMN much stronger than its physiological ligand R1810me2s-POLR2A, and our cellular studies still display that these SMN antagonists prevent SMN interaction with R1810me2s-POLR2A, resulting in the over-accumulation of active RNAP II and R-loop, mimicking depletion of *SMN*. These small molecule compounds specifically compete with methylated arginine for the binding pocket of SMN. Application of these small molecules has the advantage of maintaining the normal cellular SMN levels without disrupting methylarginine independent functions of SMN. Thus, in addition to the antisense, RNAi and CRISPR/Cas9 techniques, these potent SMN antagonists may be used as efficient tools in the study of SMN biology and its related neurological diseases.

## Online Methods

### Protein expression and purification

The coding DNA fragments of following Tudor domains were cloned into pET28-MHL vector: SMN (aa 82-147), UHRF1 (aa 126-285), SMNDC1 (aa 53-130), TDRD3 (aa 554-611), SND1 (aa 650-910), TDRD2 (aa 327-420), FXR1 (aa 2-132), PHF1 (aa 28-87), SGF29 (aa 115-293), JMJD2A (aa 897-1101), 53BP1 (aa 1483-1606), SETDB1 (aa 190-410), LBR (aa 1-67), ZGPAT (aa 120-271). The coding regions of chromodomain of CBX7 (aa 8-62), PWWP domain of DNMT3A (aa 275-417), WD40 repeats of WDR5 (aa 24-334) and CW domain of ZCWPW2 (aa 21-78) were also subcloned into pET28-MHL vector to generate N-terminally His-tagged fusion protein. The MBT repeats of L3MBTL1 (aa 200-522) and L3MBTL2 (aa 170-625) were subcloned into pET28GST-LIC vector to generate N-terminally GST-His-tagged fusion protein. The recombinant proteins were overexpressed in *E*. *coli* BL21 (DE3) Codon plus RIL (Stratagene) at 15 °C and purified by affinity chromatography on Ni-nitrilotriacetate resin (Qiagen, or Nanjing Qingning Bio-Technology Co., Ltd.) followed by TEV or thrombin protease treatment to remove the tag. The proteins were further purified by Superdex75 or Superdex200 gel-filtration column (GE Healthcare). For crystallization experiments, purified proteins were concentrated to 18 mg/mL for SMN, 23 mg/mL for UHRF1 and 10 mg/mL for TDRD3 in a buffer containing 20 mM Tris-HCl, pH 7.5, 150 mM NaCl and 1 mM DTT. All the mutations were introduced with the QuikChange II XL site-directed mutagenesis kit (Stratagene, 200522) and confirmed by DNA sequencing. Mutant proteins were also expressed in *E. coli* BL21 (DE3) Codon plus RIL and purified using the same procedures described above. The molecular weight of all protein samples was measured by mass spectrometry.

For mammalian expression, the coding DNAs of full-length SMN, SND1 and TDRD3 were cloned into mCherry2-C1 or GFP-C1 vector by restriction endonucleases *Hind* III/*Bam*H I and T4 ligase. All the mutations of full-length SMN were introduced with the QuikChange II XL site-directed mutagenesis kit (Stratagene, 200522) and confirmed by DNA sequencing.

### Small molecule fragment based screening of UHRF1 tandem Tudor domain

A small molecule fragment library with 2040 compounds was screened against tandem Tudor domain of UHRF1 by fluorescein polarization-based peptide displacement assay according to previous reports^27^. Briefly, the screening was performed in 10 μL at a protein concentration of 20 μM premixed with a 40 nM FITC-labeled H3K9me3 peptide (aa 1-25, Tufts University Core Services), and then adding a single concentration of 2 mM compound in a buffer of 20 mM Tris-HCl, pH 7.5, 150 mM NaCl, 1 mM DTT, and 0.01% Triton X-100. The hits were further confirmed by dose response analysis. All the assays were performed in duplicate in 384-well plates, using the Synergy 2 microplate reader (BioTek), with an excitation wavelength of 485 nm and an emission wavelength of 528 nm. All the compounds were purchased from Sigma or Specs company.

### Isothermal titration calorimetry (ITC)

For the ITC measurement, the concentrated proteins were diluted into 20 mM Tris-HCl, pH 7.5, 150 mM NaCl (ITC buffer); the lyophilized compounds were dissolved in the same buffer, and the pH value was adjusted by adding 2 M NaOH or 2 M HCl. The compounds that could not be dissolved in the ITC buffer were dissolved in DMSO with the accessible highest concentration. Compound concentrations were calculated from the mass and the volume of the solvent. For the ITC assay with compound dissolved in DMSO, the protein was diluted by ITC buffer containing same final concentration of DMSO. All measurements were performed in duplicate at 25 °C, using a VP-ITC (MicroCal, Inc.), an iTC-200 (MicroCal, Inc.) or a Nano-ITC (TA, Inc.) microcalorimeter. The protein with a concentration of 50-100 μM was placed in the cell chamber, and the compounds with a concentration of 0.5-2 mM in syringe was injected in 25 or 20 successive injections with a spacing of 180 s for VP-ITC or 150 or 120 s for iTC-200 or Nano-ITC. iTC-200 or Nano-ITC data should be consistent with those from the VP-ITC instrument, based on ITC results of SMN-compound **1** detected by using all the instruments. Control experiments were performed under identical conditions to determine the heat signals that arise from injection of the compounds into the buffer. Data were fitted using the single-site binding model within the Origin software 7.0 package (MicroCal, Inc.) or the independent model within the Nano-Analyze software package (TA, Inc.).

### Protein crystallization

For the complex crystal of SMN-compound **1**, compound free crystals were crystallized in a buffer containing 2 M ammonium sulfate, 0.2 M potassium/sodium tartrate, 0.1 M sodium citrate, pH 5.6 and soaked with compound **1** at a molar ratio of 1:5 for 24 h. For the complex crystals of UHRF1-compound **1**, SMN-compound **4**/**6** and TDRD3-compound **1**, purified proteins were mixed with the compounds at a molar ratio of 1:5 and crystallized using the sitting drop vapor diffusion method at 18 °C by mixing 0.5 μL of the protein with 0.5 μL of the reservoir solution. The complex of UHRF1-compound **1** was crystallized in a buffer containing 20% PEG 3350, 0.2 M magnesium nitrate; SMN-compound **4** was crystallized in a buffer containing 1.8 M sodium acetate, pH 7.0, 0.1 M Bis-Tris propane, pH 7.0; SMN-compound **6** was crystallized in a buffer containing 2 M ammonium sulfate, 0.2 M sodium chloride, 0.1 M Hepes, pH 7.5; and TDRD3-compound **1** was crystallized in a buffer containing 1.2 M sodium citrate, 0.1 M Tris-HCl, pH 8.5. Before flash-freezing crystals in liquid nitrogen, crystals were soaked in a cryoprotectant consisting of 85% reservoir solution and 15% glycerol.

### Data collection and structure determination

The program PHASER^28^ was used for molecular replacement (MR) when needed. Models were interactively rebuilt, refined and validated using COOT^29^, REFMAC^30^ and MOLPROBITY^31, 32^ software, respectively. MarvinSketch (Chemaxon.com) was used for the calculation of some SMILES strings during preparation of small molecule geometry restraints. PDB_EXTRACT^33^ and CCTBX^34^ library were used during preparation of the crystallographic models for PDB deposition and publication. Diffraction data and model refinement statistics for the structures are displayed in Table 1. Some structure determination details for specific structures are as follows. *SMN in complex with compound **1***: Diffraction images were collected on a copper rotating anode source and initially reduced to merged intensities with DENZO/SCALEPACK^35^/AIMLESS^36^. For later refinement steps, data were reduced with XDS^37^/AIMLESS. The crystal structure was solved by placement of atomic coordinates from isomorphous PDB entry 1MHN^38^ in the asymmetric unit. Geometry restraints for compound **1** were prepared on the GRADE server^39, 40^. *SMN in complex with compound **4***: Diffraction data were collected at APS/NE-CAT beam line 24-ID-E and reduced with XDS/AIMLESS. The structure was solved by MR with diffraction data from an additional, isomorphous crystal and coordinates from PDB entry 4QQ6 (SMN in complex with compound **1**, above). Geometry restraints for compound **4** were prepared with PRODRG^41^. Anisotropic displacement parameters were analyzed on the PARVATI server^42^. *SMN in complex with compound **6***: Diffraction data were collected on a rotating copper anode source and reduced with XDS/AIMLESS. The structure was solved by MR with coordinates from PDB entry 4QQ6. Geometry restraints for compound **6** were prepared with ELBOW^43^, which in turn used MOGUL. *UHRF1 in complex with compound **1***: Diffraction data were collected at APS/SBC-CAT beamline 19ID and reduced to merged intensities with XDS/AIMLESS. The structure was solved by MR with coordinates derived from PDB entry 3DB3^44^. *TDRD3 in complex with compound **1***: Diffraction data were collected at CLS/CMCF beamline 08ID and reduced to intensities with DENZO/SCALEPACK. Intensities were converted to the MTZ format with COMBAT^45^ or, alternatively, POINTLESS^46^ before symmetry-related intensities were merged with AIMLESS. The structure was solved by MR with coordinates from PDB entry 3PMT^18^.

### Fluorescence analysis

U2OS cells were plated in a 35 mm FluoroDish with a 0.17 mm coverslip bottom (World Precision Instruments, FD35-100) for 12∼24 h and transfected with 1.0 μg mCherry-SMN WT (wild-type) or mutant plasmids. Media was changed 4∼6 h after transfection and cells were cultured for another 24 h. 10 μM CCVJ-Cmpd **1** was added for 24 h treatment. Then the treated cells were rinsed and the media replaced with phenol-free FluoroBright DMEM for analysis by using Zeiss LSM880 microscopy.

### Pulldown and western blotting

U2OS cells were lysed in RIPA buffer (140 mM NaCl, 10 mM Tris-HCl, pH 7.6, 1% Triton, 0.1% sodium deoxycholate, 1 mM EDTA) containing protease inhibitors (Roche, 05892791001). The cell lysate was incubated with 20 μM biotin-Cmpd **1** overnight at 4 °C, then 30 μL streptavidin beads (ThermoFisher Scientific, 20353) was added and incubated at 4 °C for 1 h. The beads were then washed with RIPA buffer for 3 times, and loading buffer was added and boiled for 5 min for elution. The eluted samples were loaded on SDS-PAGE gel for western blotting (SMN antibody, BD Transduction Laboratories, 610646; TDRD3 antibody, Cell signaling, 5492; SND1 antibody, Bethyl, A302-883A; Streptavidin-Horseradish Peroxidase (HRP), Invitrogen, SA10001; RFP antibody, Abcam, ab62341; GFP antibody, Santa Cruz Biotechnology, sc-9996) or mass spectrometry (MS) analysis. For the competition analysis, 100 μM biotin-free compound **1** or compound **15** was added at the same time when the biotin-Cmpd **1** was incubated with the cell lysates.

### Affinity-purification and mass spectrometry (AP-MS)

The samples for AP-MS were prepared following pulldown procedure as previously described^47^. Briefly, the eluted samples were loaded on NuPAGE gel (Invitrogen, NP0326BOX), the gels with samples contained were cut and submitted to Proteomics Facility in University of Texas at Austin for protein identification analysis. The samples were digested, desalted run on the Dionex LC and Orbitrap Fusion 2 for LC-MS/MS with a 2-hour run time and processed by using PD 2.2 and Scaffold 5. The data of AP-MS were analyzed as previously described method with minor modifications^48^. Briefly, the peptide spectrum matches (PSMs) of the identified proteins were extracted from AP-MS datasets for the experimental sample (equal quantity DMSO added), biotin-free compound **1** competitive sample and biotin-free compound **15** non-competitive sample, respectively. The relative protein intensity ratios were obtained by PSM of experimental sample to competitive sample, PSM_Expt_/PSM_Cmpt_, and PSM of non-competitive sample to competitive sample, PSM_NonCmpt_/PSM_Cmpt_, respectively. If the PSM was equal to zero, 0.5 was assigned in order to avoid yielding infinite value after dividing zero during the data processing. The normalized PSM_Expt_/PSM_Cmpt_ was plotted along with normalized PSM_NonCmpt_/PSM_Cmpt_ by using Origin software 7.0. Proteins with an intensity ratio > 4 from both PSM_Expt_/PSM_Cmpt_ and PSM_NonCmpt_/PSM_Cmpt_ were considered as biotin-Cmpd **1** enriched proteins.

### Cell culture

HEK293 were grown in DMEM (SLRI media facility) plus 10% FBS (Sigma, F1051). For analysis of SMN chemical antagonists, HEK293 cells were treated with DMSO or a serious of concentrations (0, 2, 6, 10, 20, 30, 40 and 80 μM) of compound **1**, compound **2** or negative compound **15** for 72 h. CRISPR-mediated *SMN1* gene knockout was performed according to our previous study^9^. Briefly, 2 μg of CRISPR/Cas9 plasmid (pCMV-Cas9-GFP), which expresses scrambled guide RNA, or guide RNA that targets the *SMN1* gene exon1 (gRNA target sequence: ATTCCGTGCTGTTCCGGCGCGG) or exon3 (gRNA target sequence: GTGACATTTGTGAAACTTCGGG) was transfected into HEK293 cells. Cells were sorted by BD FACSAria flow cytometry at Donnelly Centre, University of Toronto 24 h after transfection and single GFP-positive cells were seeded into a 48-well plate. The expression levels of SMN in each clone were detected by immunofluorescence. The transfection of GFP-RNase H1 R-loop-binding domain (GFP-HB) for R-loop detection into HEK293 cells was performed with the FuGENE Transfection reagent (Roche, E269A).

### Immunoprecipitation (IP) and western blotting

HEK293 cells were subjected to three freeze-thaw cycles in high-salt lysis buffer (10 mM Tris-HCl, pH 7.9, 10% glycerol, 420 mM NaCl, 0.1% Nonidet P-40, 2 mM EDTA, 2 mM DTT, 10 mM NaF, 0.25 mM Na_3_VO_4_, and 1× protease inhibitor mixture (Sigma, P8340)), followed by centrifugation at 14,000 rpm for 1 h at 4 °C to remove insoluble materials. The supernatant cell lysates were sonicated with five on and off cycles of 0.3 s/0.7 s per 1 mL and incubated for 30 min at 4 °C with 12.5 –25 units/mL benzonase nuclease (Sigma, E1014) to remove RNA and DNA, followed by centrifugation at 14,000 rpm for 30 min at 4 °C. The supernatant cell lysates were incubated with 2 μg of antibody overnight at 4 °C, followed by addition of 20 μL of Dynabeads Protein G beads (Invitrogen, 10004D) for an additional incubation for 4 h. After washing with low-salt buffer (10 mM Tris-HCl, pH 7.9, 100 mM NaCl, and 0.1% Nonidet P-40), associated proteins were eluted into protein-loading buffer and separated by Tris 4–20% SDS-polyacrylamide (Mini-PROTEAN TGX Precast Protein Gel, BioRad, 4561096), and transferred to nitrocellulose or PVDF membranes (Immu-Blot PVDF, BioRad, 1620112 or 1620177). Transferred samples were immunoblotted with primary antibodies (POLR2A, Abcam, ab5408; SMN, Santa Cruz Biotechnology, sc-15320; ACTB, Sigma, A5441; GFP, Invitrogen, G10362; TUBB, Santa Cruz Biotechnology, sc-9104) at a dilution of 1:2,000 to 1:5,000, followed by horseradish peroxidase-conjugated goat anti-mouse or mouse anti-rabbit secondary antibody (Jackson Immuno Research, 211-032-171 or 115-035-174) at a dilution of 1:10,000. Western blot detection was performed with enhanced chemiluminescence (Pierce ECL Western Blotting Substrate, Thermo Scientific, 32209). For analysis of SMN chemical antagonists, HEK293 cells were treated with DMSO, compound **1** or compound **2** with a series of concentrations for 72 h, before being processed for IP and western blotting.

### Chromatin immunoprecipitation (ChIP)

ChIP was performed using the EZ-ChIP™ A - Chromatin Immunoprecipitation Kit (Millipore, 17-371) according to the manufacturer’s instruction. Antibodies were used with a range of 1-2 μg, and IgG (Millipore, polyclonal antibody, 12-370) was used as a background control. After immunoprecipitation, genomic DNA was de-crosslinked in ChIP elution buffer containing 5 M NaCl at 65 °C overnight and purified with t he Qiaex II kit (Qiagen, 20021) and eluted in water for PCR amplification. Immunoprecipitated and input DNAs were used as templates for qPCR. The qPCR primer sequences for *ACTB* gene are the same as described earlier^9^. For analysis of SMN chemical antagonists, HEK293 cells were treated with DMSO, compound **1** or compound **2** (a final concentration of 20 μM) for 72 h, before being processed for ChIP.

### Immunofluorescence and microscopic R-loop quantification

Global nuclear R-loop detection was ascertained via R-loop immunofluorescence using S9.6 antibody for R-loop (Kerafast, ENH001). 24 h prior to immunofluorescence, 40,000 cells were seeded on to Poly-L-Lysine (PLL) coated coverslips. Cells were fixed using 1% formaldehyde for 15 min, washed three times with PBS, permeabilized with 0.3% Triton X-100 and then washed again three times with PBS. Coverslips were blocked using 5% BSA for 1 h at room temperature and transferred to humidified chambers for antibody incubations. Coverslips were incubated with 60 μL of S9.6 (1:500 dilution) for 1 h at room temperature. After washing with PBS, cells were incubated with secondary antibody for 1 h in a dark chamber. Following further washing and DAPI staining, coverslips were mounted onto microscope slides using DAKO fluorescent mounting medium and then sealed with nail polish. For RNaseH1 overexpression analysis, scramble and SMN knockout cells were seeded in 6-well plates and transfected 24 h later with 0.9 μg pcDNA3-Empty or pcDNA3-RNaseH1. At 48 h post-transfection, cells were harvested and re-seeded onto PLL-coated coverslips, which were processed for immunofluorescence 24 h later using S9.6 and anti-RNaseH1 (Proteintech, 15606-1-AP) to quantify R-loop and confirm RNaseH1 overexpression, respectively. For analysis of SMN chemical antagonists, HEK293 cells were treated with DMSO, compound **1** or compound **2** (a final concentration of 20 μM) for 72 h, before being processed for S9.6 immunofluorescence.

We employed a Nikon C2+ confocal microscope coupled to NIS-elements AR software (Nikon). For R-loop microscopy in HEK293 cells, random fields identified by DAPI staining were captured at 100× magnification. For any given image, 5-6 2D imaging planes were acquired along the z-axis to generate 3D confocal image stacks. DAPI was used to create masks of nuclei and S9.6 intensity values for individual cells were obtained as maximum intensity planes via the NIS-elements AR software (Nikon). Representative single-plane images from z-stacks adjusted for background and contrast in Photoshop (Adobe) are shown.

### Data availability

Coordinates and structure factors are deposited in the Protein Data Bank (PDB) with accession codes 4QQ6, 4QQD, 7W2P, 7W30, 6V9T for SMN-compound **1**, UHRF1-compound **1**, SMN-compound **4**, SMN-compound **6**, TDRD3-compound **1**, respectively. All other relevant data supporting the key findings of this study are available within the article or from the corresponding authors upon reasonable request.

## Acknowledgments

We thank Dr. Wolfram Tempel for data collection and structure determination, Dr. John R. Walker for reviewing some of the crystal structures, and Dr. Dalia Barsyte-Lovejoy, Dr. Magdalena M Szewczyk and Dr. Hui Peng for technical assistance. The Structural Genomics Consortium is a registered charity (no: 1097737) that receives funds from Bayer AG, Boehringer Ingelheim, Bristol Myers Squibb, Genentech, Genome Canada through Ontario Genomics Institute [OGI-196], EU/EFPIA/OICR/McGill/KTH/Diamond Innovative Medicines Initiative 2 Joint Undertaking [EUbOPEN grant 875510], Janssen, Merck KGaA (aka EMD in Canada and US), Pfizer and Takeda. This work was also supported by a NSERC grant RGPIN-2021-02728 (J.M.); the National Natural Science Foundation of China grants (31500615 (Y.L.) and 81773608 (X.J.)); the Priority Academic Program Development of the Jiangsu Higher Education Institutes (PAPD); and CPRIT grant RP180804 and the NIH grant GM126421 (M.T.B.). Some diffraction experiments were performed at the Structural Biology Center and Northeastern Collaborative Access Team (NIGMS grant P30 GM124165) and Structural Biology Center beam lines at the Advanced Photon Source at Argonne National Laboratory. ANL is operated by the University of Chicago Argonne, LLC, for the U.S. Department of Energy Office of Biological and Environmental Research under contract DE-AC02-06CH11357. Diffraction experiments described in this paper were also performed by using beamline 08ID-1 at the Canadian Light Source, which is supported by the Canada Foundation for Innovation, Natural Sciences and Engineering Research Council of Canada, the University of Saskatchewan, the Government of Saskatchewan, Western Economic Diversification Canada, the National Research Council Canada, and the Canadian Institutes of Health Research. Protein identification was provided by the UT Austin Center for Biomedical Research Support Proteomics Facility, which is run by Maria D. Person.

## Author contributions

Y.L. and W.L. purified and crystallized the proteins; A.I. conducted the fragment based screening under the supervision of P.B.; Y.L., W.L., S.Q., M.Z. and M.Z. conducted the ITC assays; Z.N., D.Y.Z, K.J.A., and X.G. performed the cellular assay under the supervision of K.M. and J.G.; Y.W. performed the fluorescence analysis and pulldown assay under the supervision of M.T.B.; J.R. synthesized the modified compounds under the supervision of X.J.; P.L. and Y.L. cloned the constructs; X.Z. and G.X. analyzed the MS data and made the associated figures; Y.L. and J.M. conceived the study and wrote the paper with substantial contributions from all the other authors.

## Competing financial interests

The authors declare no competing financial interests.

## Additional methods

Synthesis of CCVJ and biotin conjugated compound **1** (CCVJ-Cmpd **1** and biotin-Cmpd **1**) and the respective characterization are reported in the Supplementary Note.

---

**Extended Data Fig. 1.**
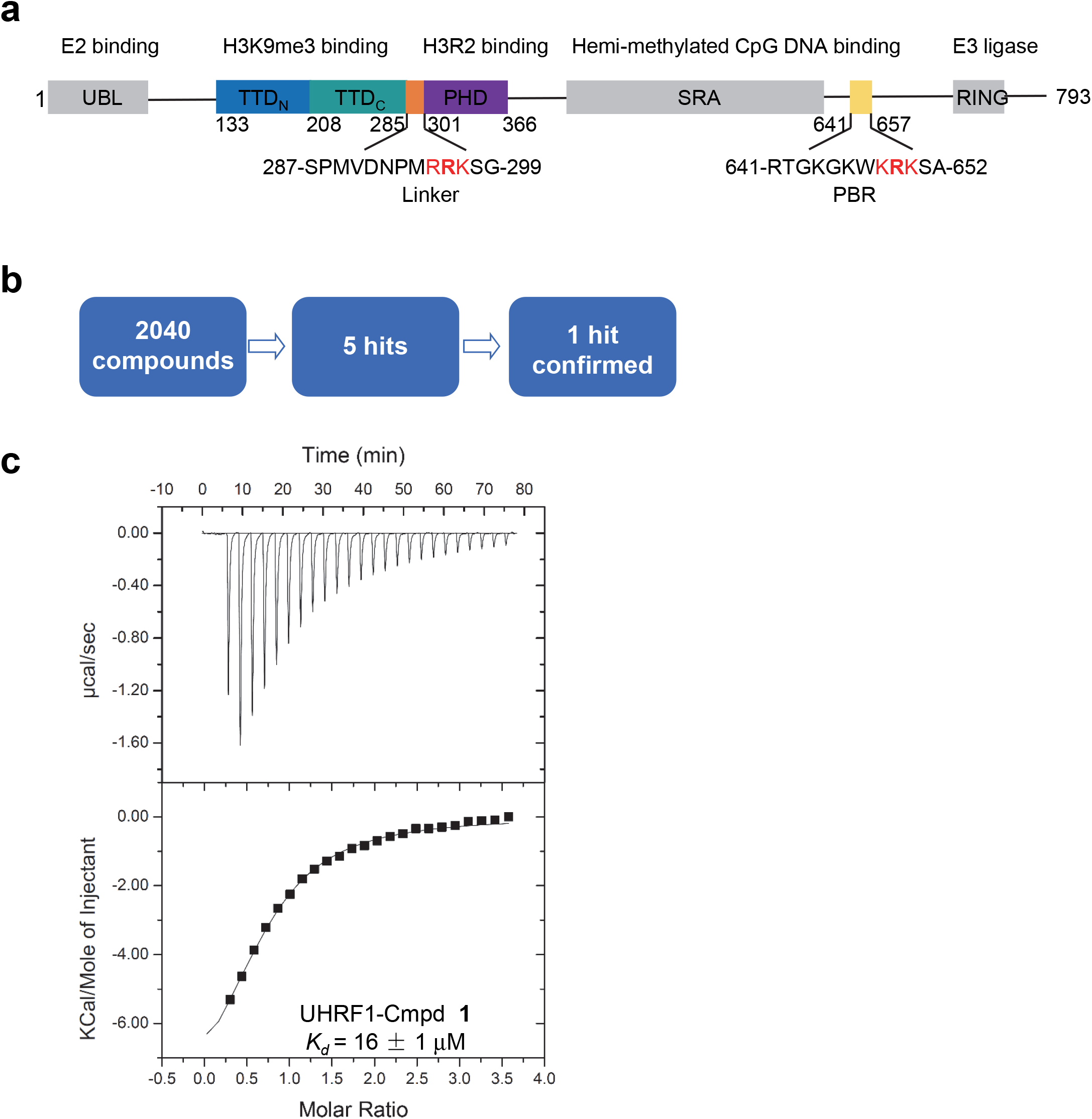
Fragment screening for tandem Tudor domain (TTD) of UHRF1. **a**, Domain structure of UHRF1. UBL, ubiquitin-like domain; TTD, tandem Tudor domain, containing TTD_N_ and TTD_C_ sub-domains; PHD, plant homeodomain; SRA, SET and RING associated domain; PBR, polybasic region; RING, really interesting new gene. **b**, Flow-chart of fragment screening for TTD of UHRF1. **c**, ITC binding curve for the titration of compound **1** to TTD of UHRF1. ITC data shown are representative of two independent experiments.

**Extended Data Fig. 2.**
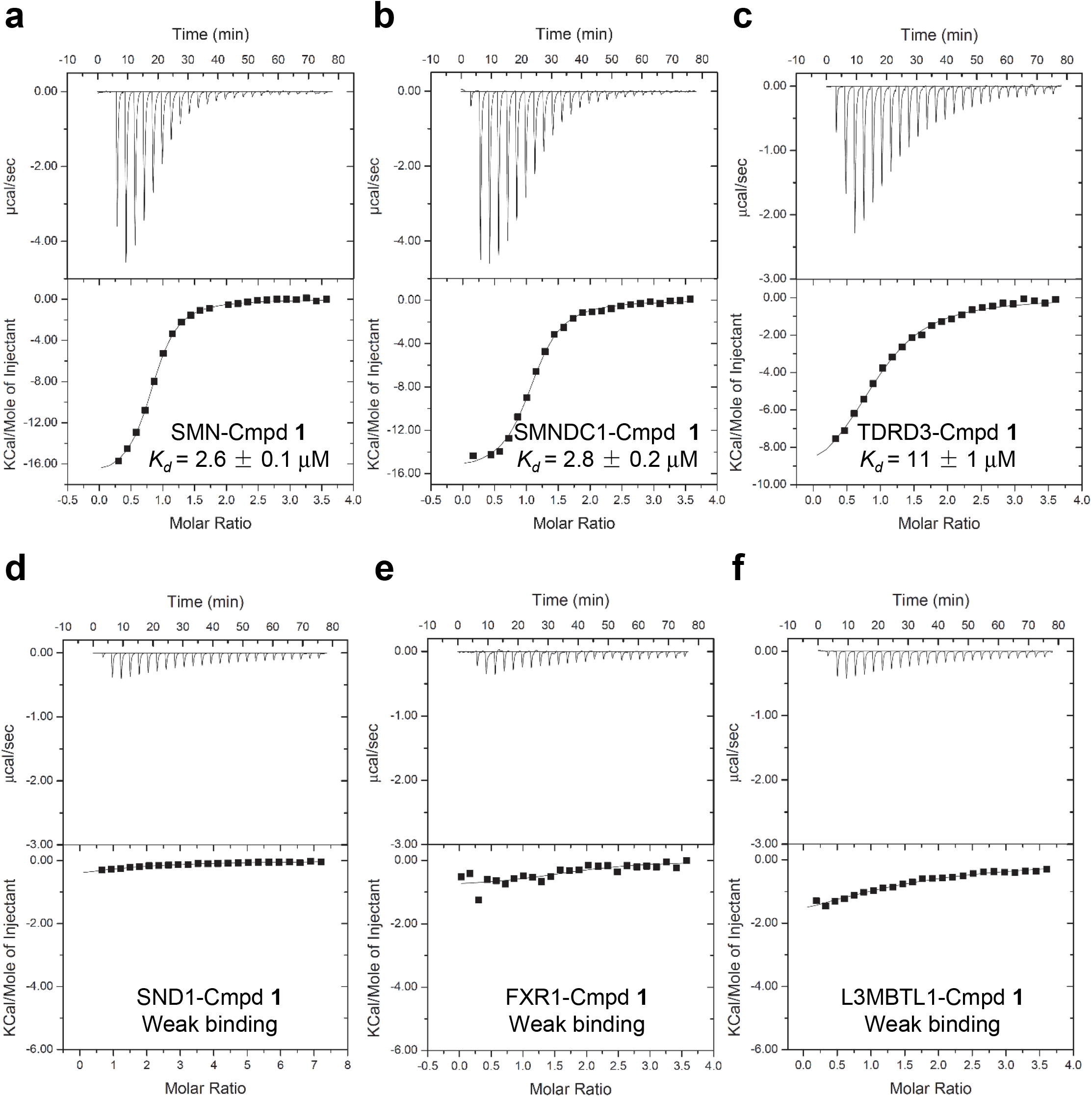
Compound 1 prefers to binding to Tudor domain of SMN. ITC binding curves for the titration of compound **1** to the Tudor domain of **a**, SMN, **b**, SMNDC1, **c**, TDRD3, **d**, SND1, **e**, FXR1, and **f**, MBT repeats of L3MBTL1, respectively. ITC data shown are representative of two independent experiments.

**Extended Data Fig. 3.**
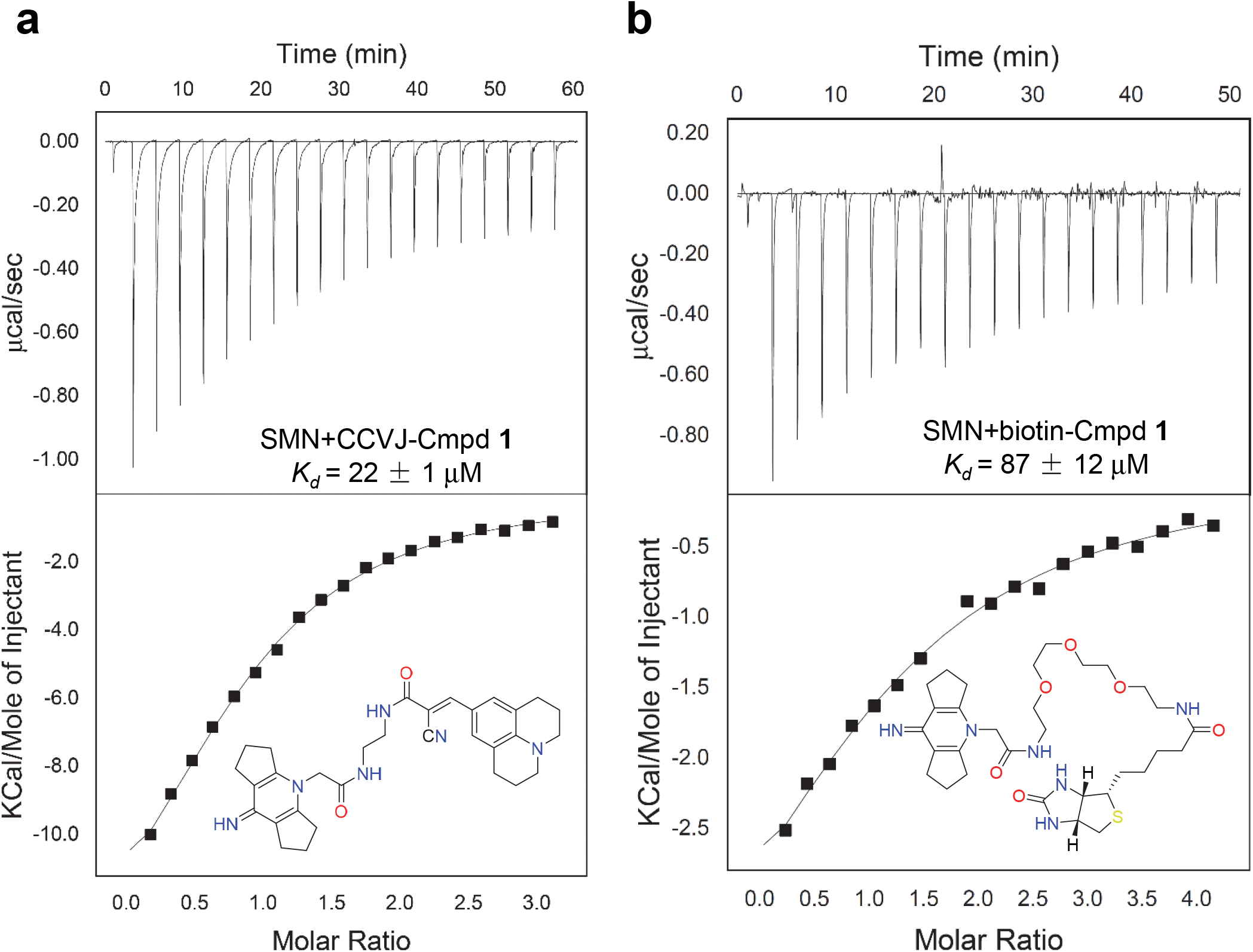
Modified compound 1s still bind to Tudor domain of SMN. ITC binding curves for the titration of CCVJ-Cmpd **1** (**a**) or biotin-Cmpd **1** (**b)** to the Tudor domain of SMN, respectively. ITC data shown are representative of two independent experiments.

**Extended Data Fig. 4.**
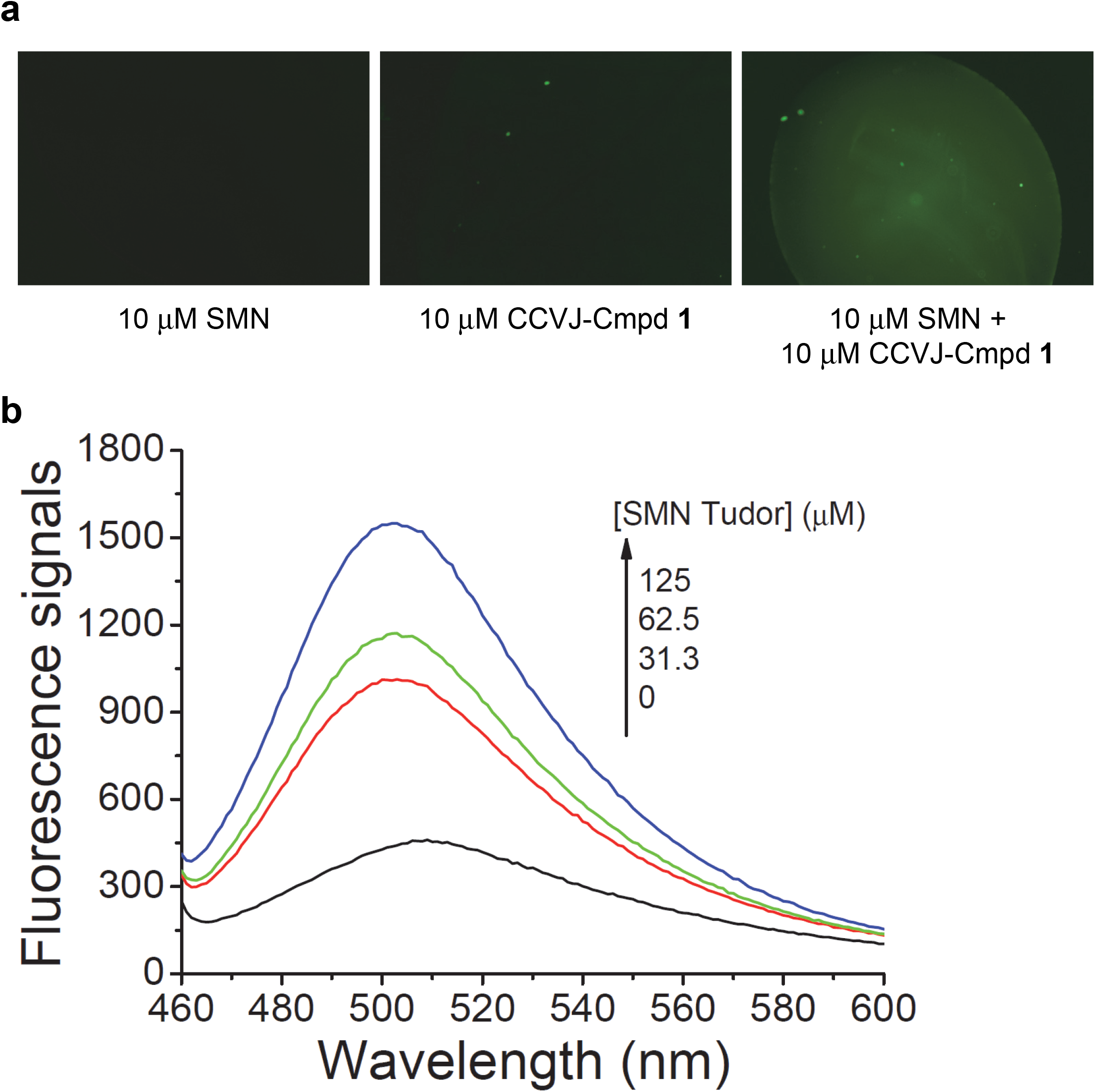
Interaction between Tudor domain of SMN and CCVJ-Cmpd 1 triggers fluorescence signals. **a**, Fluorescence signals of 10 μM SMN Tudor, 10 μM CCVJ-Cmpd **1**, and mixture of 10 μM SMN Tudor with 10 μM CCVJ-Cmpd **1** were detected by fluorescence microscope (Nikon, ECLIPSE Ti-E), under the excitation of blue light. The CCVJ-Cmpd **1** only showed weak fluorescence signal, which was enhanced by addition of the SMN Tudor domain protein. Exposure time: 1 s. **b**, Fluorescence signals for addition of different concentrations of the SMN Tudor domain protein to 5 μM CCVJ-Cmpd **1** detected by Infinite M1000 Pro microplate reader (Tecan) with an excitation wavelength of 440 nm and an emission wavelength of 460-600 nm. Data shown are representative of three independent experiments.

**Extended Data Fig. 5.**
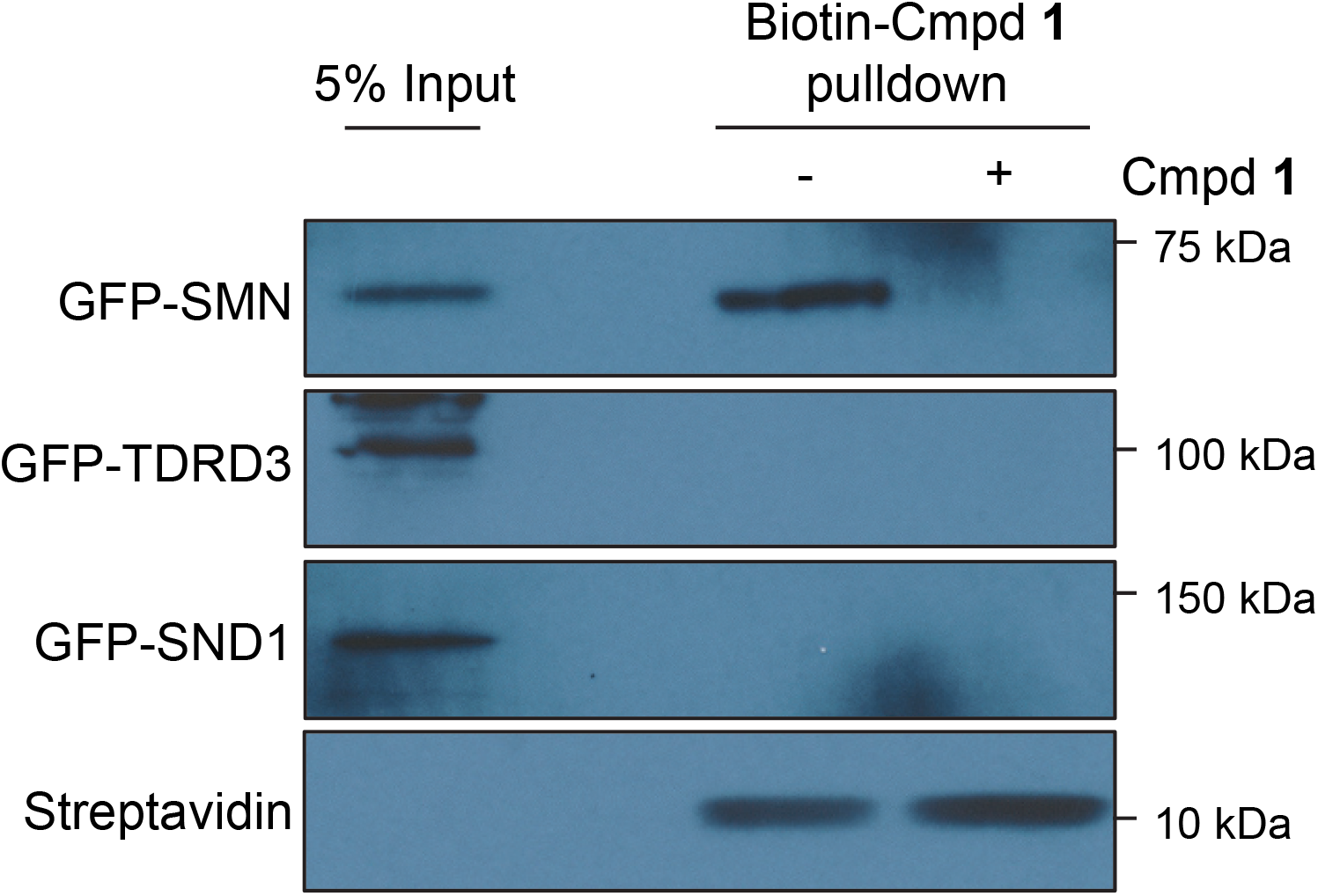
Biotin-Cmpd 1 selectively chemiprecipates SMN but not TDRD3 or SND1, another two Tudor domain-containing proteins from GFP-SMN/TDRD3/SND1 co-transfected U2OS cells. Data shown are representative of three independent experiments.

**Extended Data Fig. 6.**
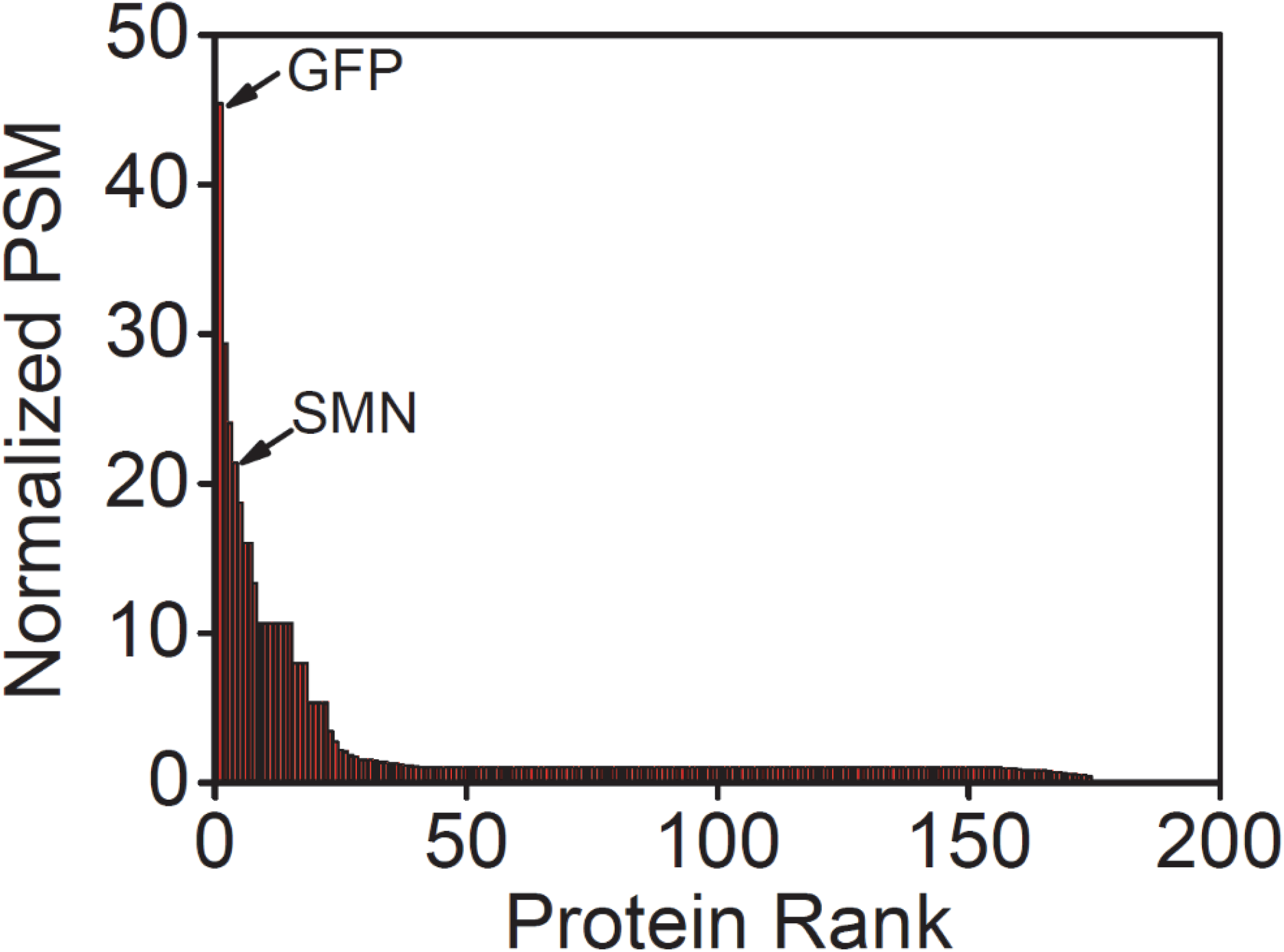
Biotin-Cmpd 1 pulldown and AP-MS analysis of GFP-SMN overexpressed U2OS cell lysate indicates specific cellular engagement of compound 1 to SMN. The GFP-SMN PSMs were normalized by GFP-EV PSMs and the final figure was drawn by Origin software 7.0. The data showed that SMN was enriched specifically by the biotin-Cmpd **1,** accompanied by two other common contaminants of the U2OS cell lysate, KRT8 (Keratin, type II cytoskeletal 8) and HNRNPL (Heterogeneous nuclear ribonucleoprotein L).

**Extended Data Fig. 7.**
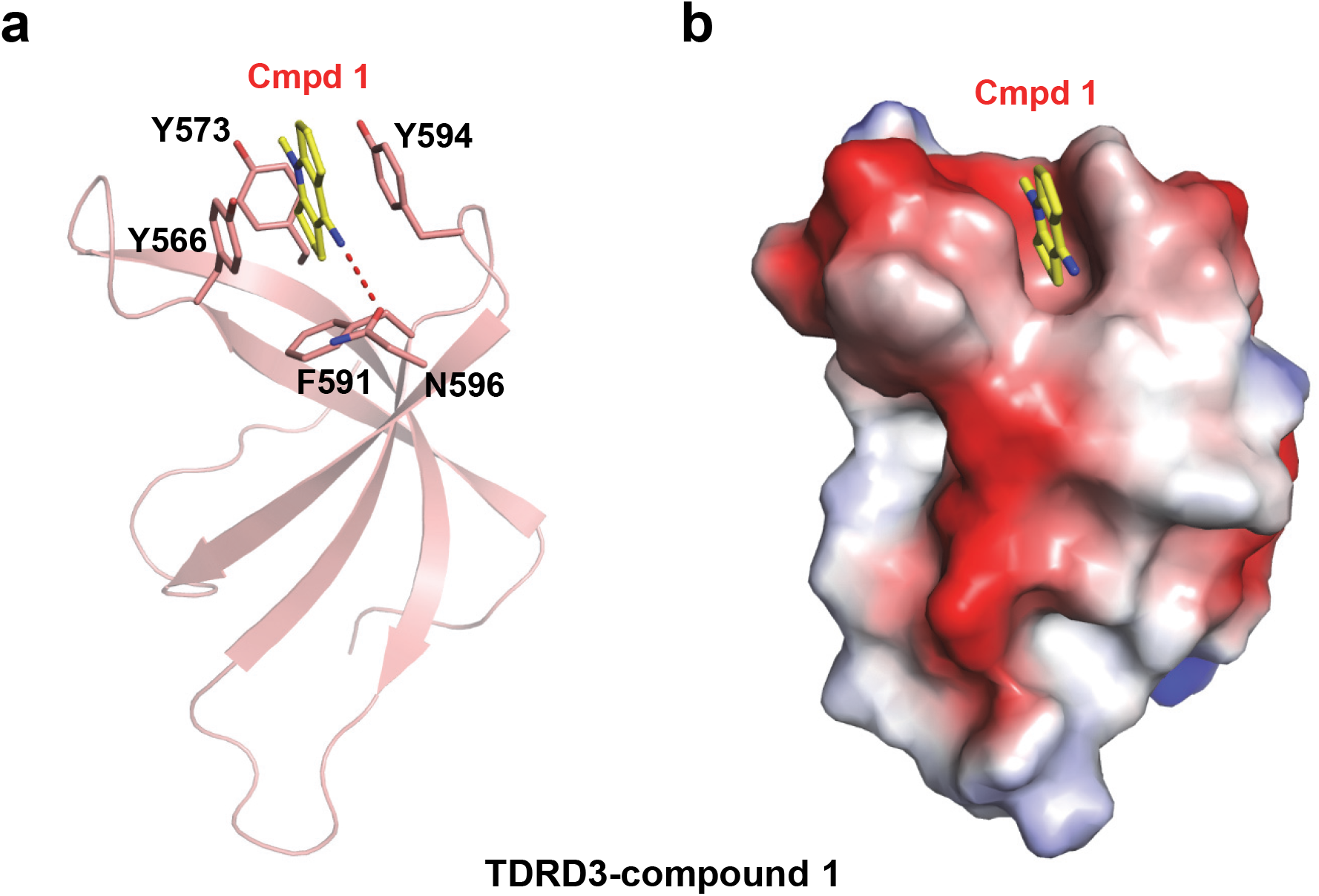
Complex structure of Tudor domain of TDRD3 and compound 1. **a**, The TDRD3 complex structure is shown in a cartoon mode. The Tudor domain of TDRD3 is colored in salmon, with the interacting residues shown in sticks and the intermolecular hydrogen bonds shown in red dashes. **b**, Electrostatic potential surface representation of the complex.

**Extended Data Fig. 8.**
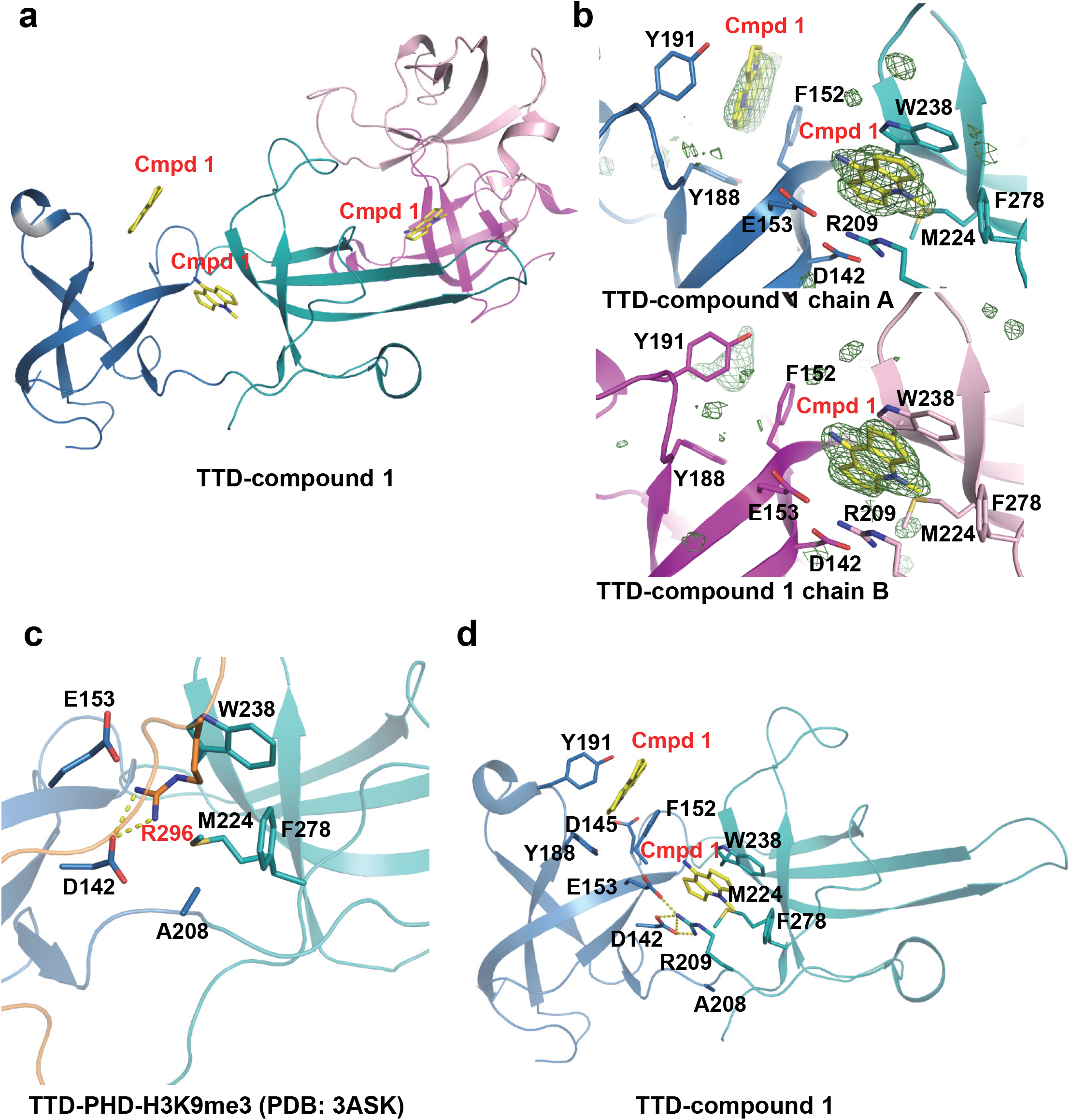
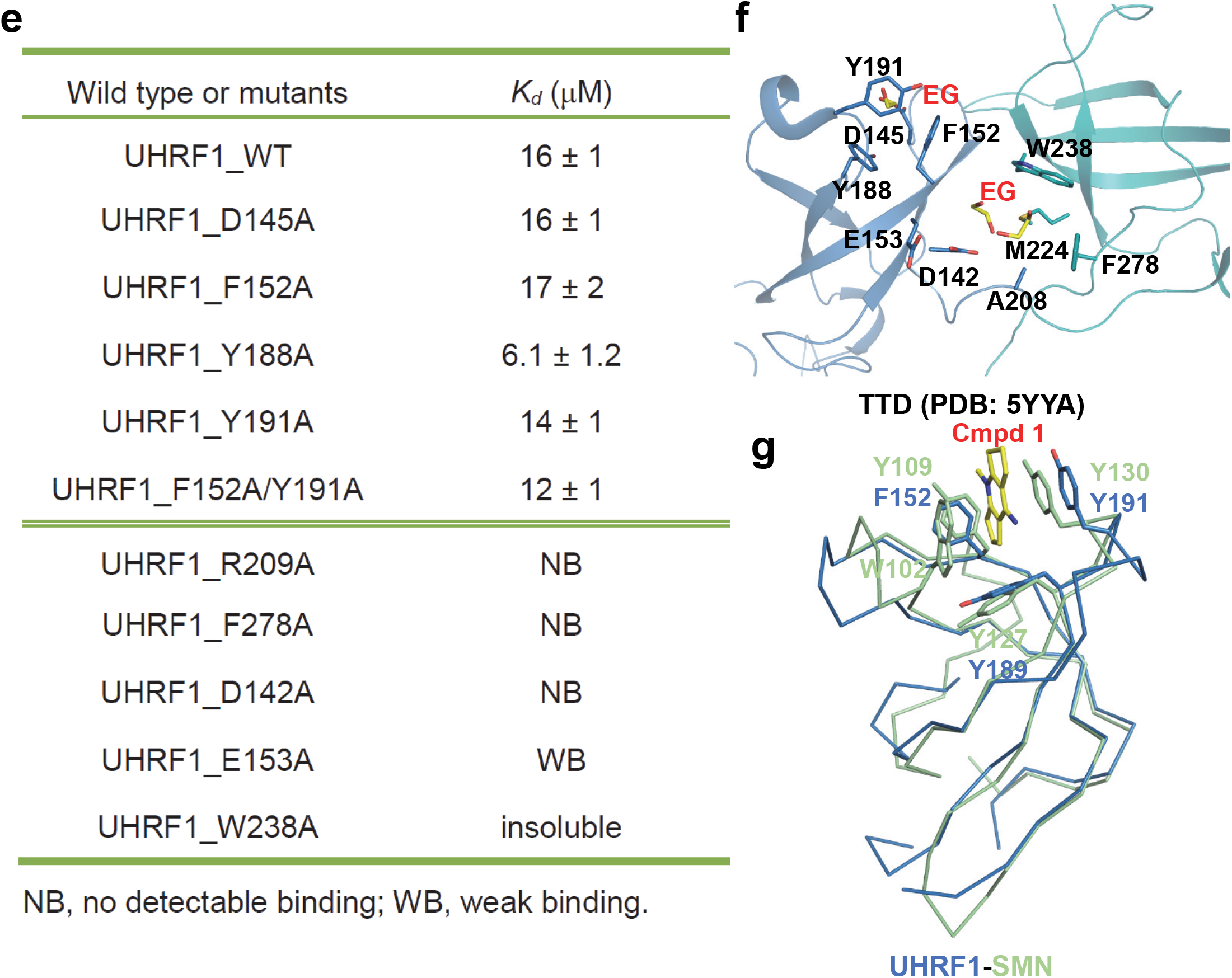
The aromatic cage of UHRF1 is not the binding site for compound 1. **a**, Overall structure of TTD domain of UHRF1 in complex with compound **1**. The TTD_N_ and TTD_C_ of the two UHRF1 molecules were colored in blue, green (chain A) and magenta, pink (chain B), respectively. **b**, Complex structure of UHRF1_TTD-compound **1** with the Fo–Fc omit map of compound **1** contoured at 3 σ and TTD shown in a ribbon mode. **c**, Complex structure of TTD-PHD of UHRF1 with an H3K9me3 peptide (PDB: 3ASK). The binding pocket of residue R296 from the TTD-PHD linker is shown in a stick mode and the intramolecular hydrogen bonds were shown in yellow dashes. **d**, Complex structure of TTD of UHRF1 and compound **1** is shown in a cartoon mode with the interacting residues shown in sticks and the intramolecular hydrogen bonds indicated by yellow dashes. **e**, Binding affinities of compound **1** to different UHRF1 TTD mutants determined by ITC. ITC data shown are representative of two independent experiments. **f**, Apo structure of the UHRF1 TTD domain with small molecules (EG, ethylene glycol from the crystallization buffer) located in the H3K9me3 and arginine binding pockets. **g**, Comparison of the methyllysine binding cage of UHRF1 to the methylarginine binding cage of SMN. The cage forming residues and compound **1** are shown in sticks.

**Extended Data Fig. 9.**
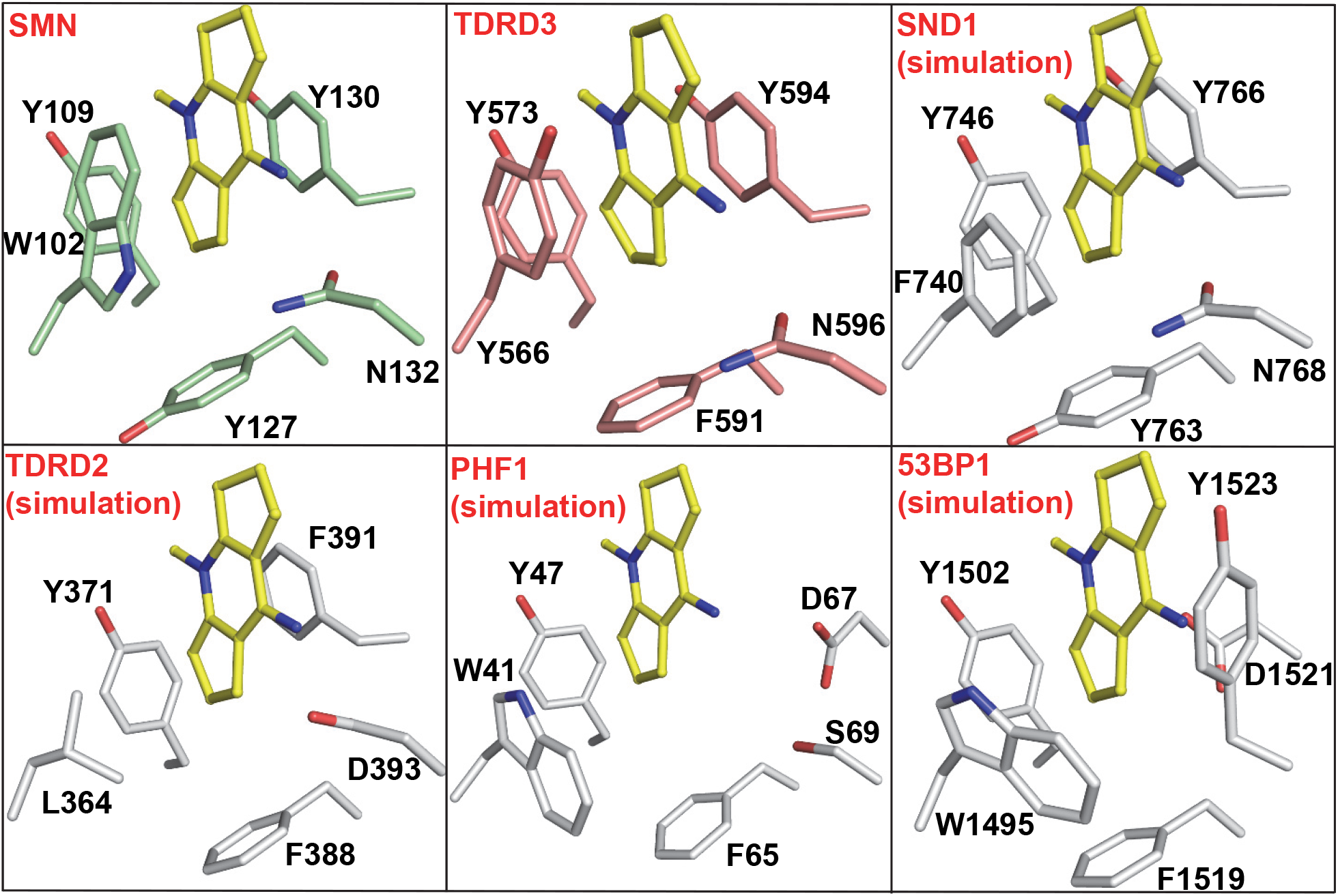
Complex structures or complex models of different Tudor domains with compound 1. The complex models were built based on superposition of different Tudor domain structures onto the SMN-compound **1** complex structure.

**Extended Data Fig. 10.**
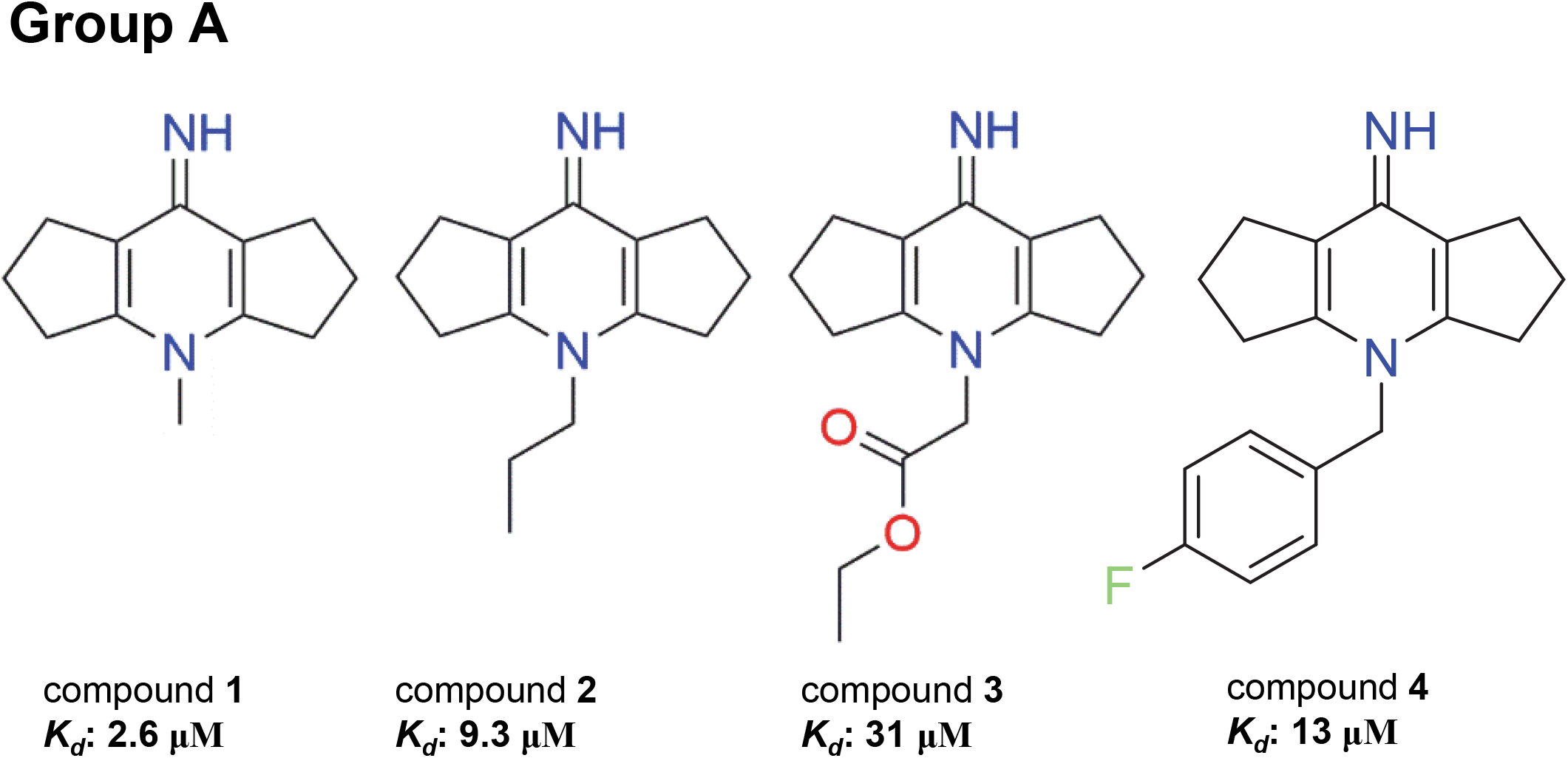
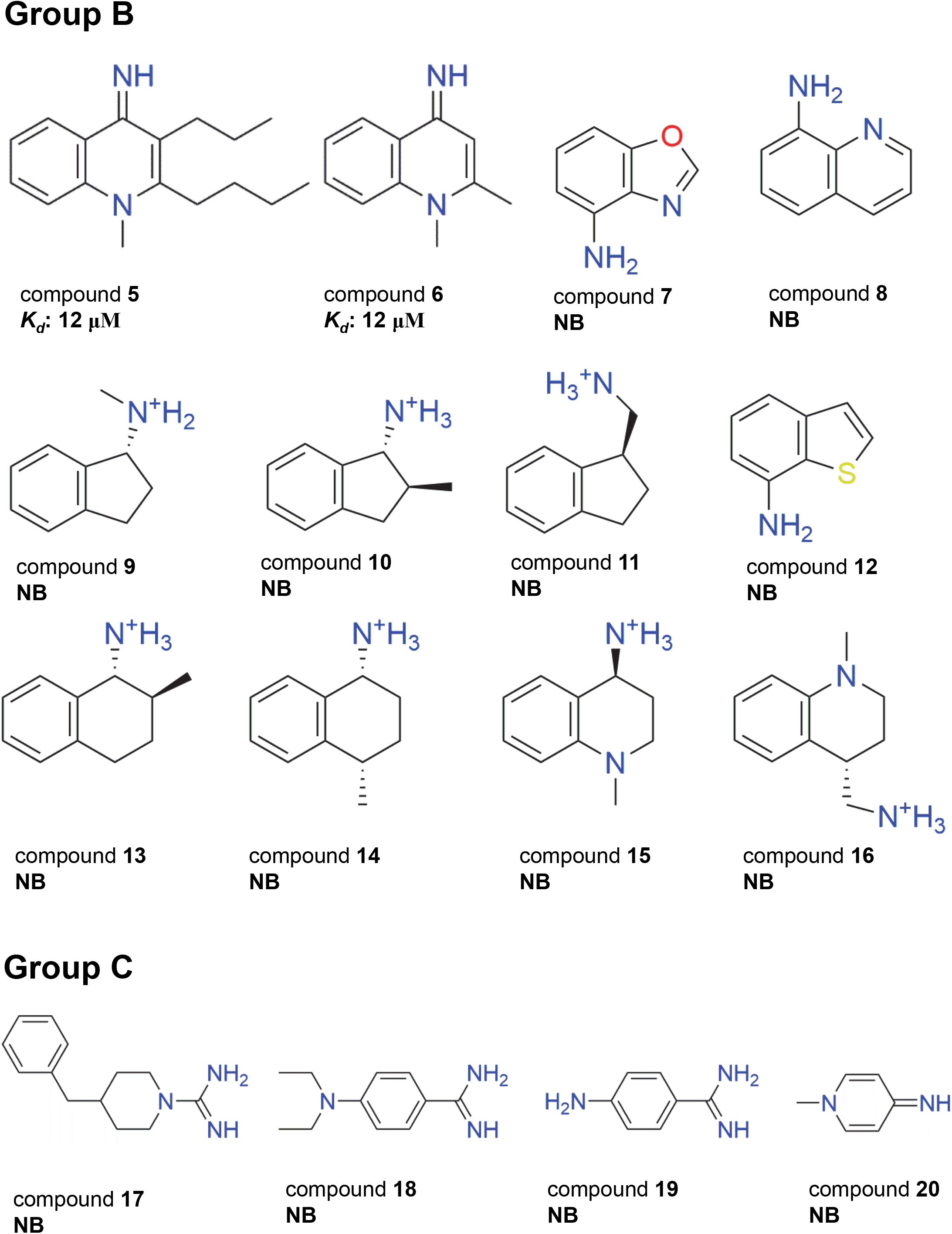
Chemical structure and binding affinity (determined by ITC) of different compounds to SMN.

**Extended Data Fig. 11.**
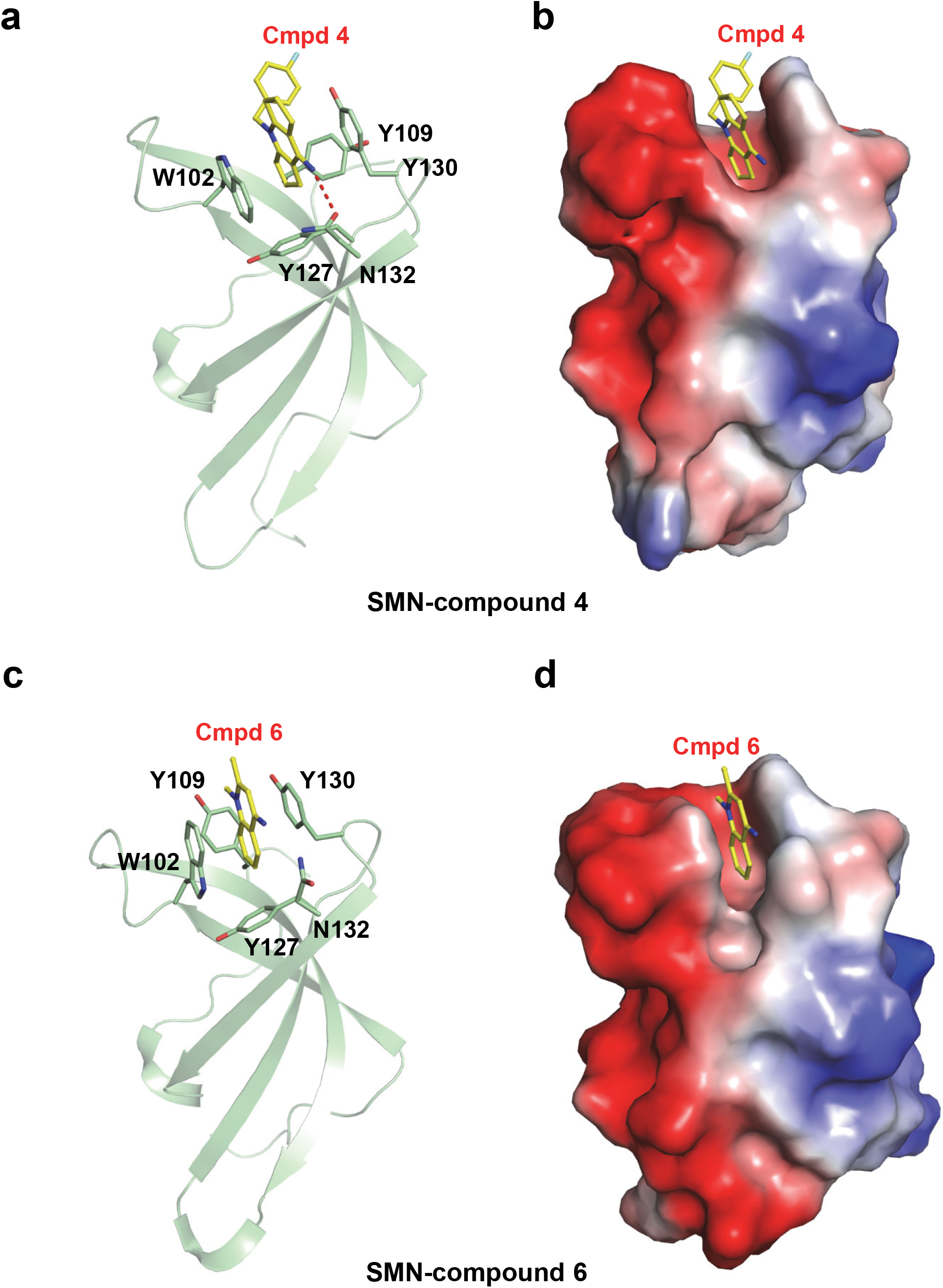
Complex structures of SMN-compound 4 and SMN-compound 6. Complex structure of Tudor domain of SMN and compound **4** or **6** shown in a cartoon mode (**a** or **c**) and electrostatic potential surface representation (**b** or **d**), respectively.

**Extended Data Fig. 12.**
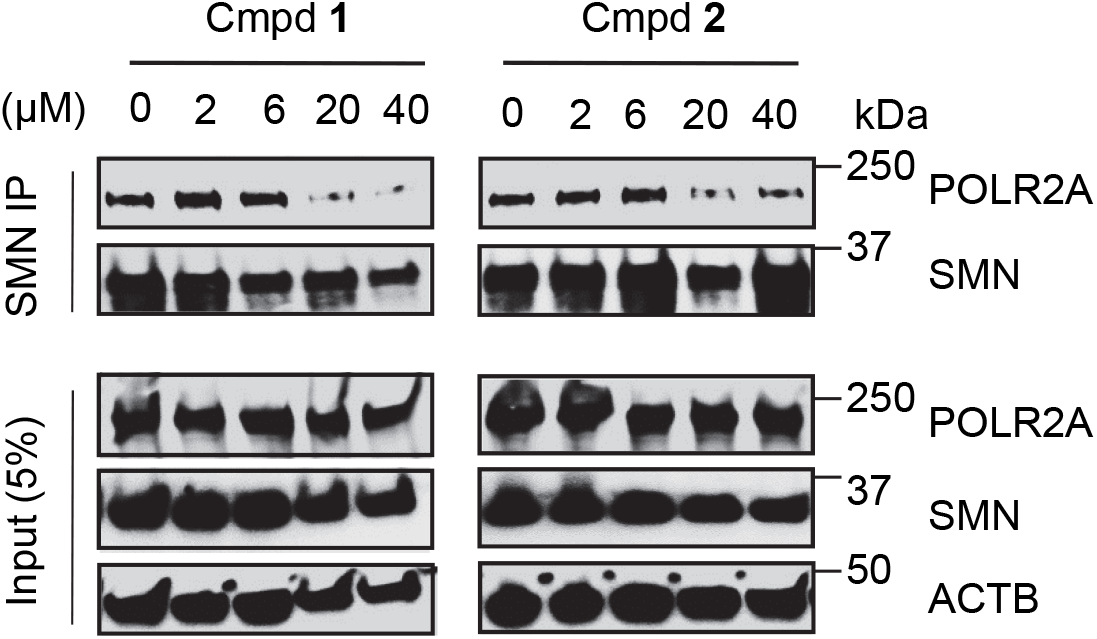
The SMN antagonists disrupt binding of SMN to RNAP II. IP-western blot experiments were performed by using the indicated antibodies in HEK293 cell extract treated with indicated concentrations of compound **1**, compound **2**, or DMSO for 72 h. Data shown are representative of three independent experiments.

**Extended Data Fig. 13.**
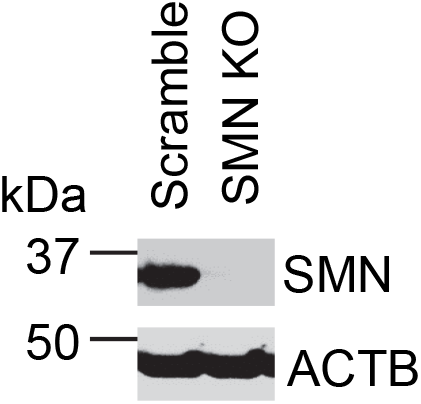
The efficiency of knockout of SMN by CRISPR/Cas9 was examined by western blot analysis. Data shown are representative of three independent experiments.

**Extended Data Fig. 14.**
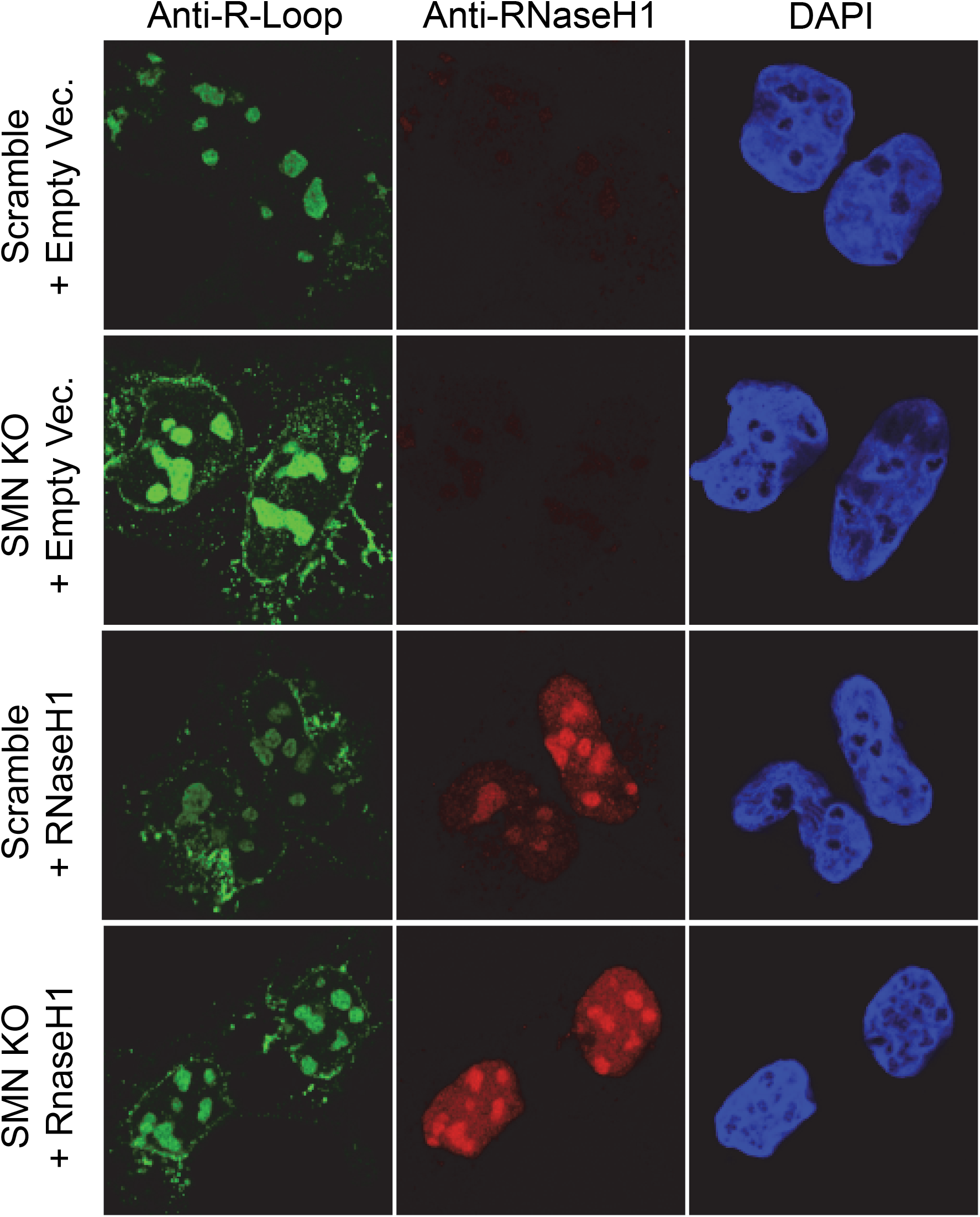
RNaseH1 overexpression decrease R-loop accumulation. Double immunofluorescence coupled to confocal microscopy analysis of R-loop staining in SMN-knockout HEK293 cells expressing human RNaseH1 or empty control vector. Shown are representative confocal images of three independent experiments.

## Notes

### Competing Interest Statement

The authors have declared no competing interest.

